# Nutrient landscape shapes the genetic diversification of the human gut commensal *Bacteroides thetaiotaomicron*

**DOI:** 10.1101/2025.06.24.661248

**Authors:** Michaela Lang, Christos Zioutis, Andreas Heberlein, Nika Ivanovova, Jasmin Schwarz, Stephan Köstlbacher, Olga Bochkareva, Kayla Jean Flanagan, Fátima C. Pereira, David Berry

## Abstract

*Bacteroides thetaiotaomicron* is a prominent member of the human gut microbiome that has evolved a suite of polysaccharide utilization loci (PUL) to break down a range of diet- and host-derived glycans. To gain insight into the evolution of this bacterium on ecologically meaningful time scales, we carried out an *in vitro* evolution study in which *B. thetaiotaomicron* was cultivated on media of different carbohydrate complexity for three months. Shotgun sequencing of the evolved populations revealed an increased number of single nucleotide polymorphisms with increased medium complexity, suggesting that genetic diversification is driven in part by the nutrient landscape. We also observed high-frequency reversible DNA inversions mediated by site-specific DNA integrases, which may be important to production and maintenance of phenotypic heterogeneity in the population. Competition experiments against the ancestor revealed adaption to the experimental conditions, with a fitness gain of the evolved populations and lineages thereof. This fitness advantage was accompanied by an increase in cell size, faster glucose depletion rates, and increased amylopectin degradation in the presence of glucose. In conclusion, rapid adaptive genetic diversification in *B. thetaiotaomicron* is induced and maintained in part by a complex nutritional environment.

## Introduction

The human gut microbiome harbors hundreds of bacterial species that are important for the degradation of dietary components that are not readily absorbed by the host. Bacteroides thetaiotaomicron is a commensal human gut bacterium that is highly specialized in the degradation of dietary and host-derived glycans [1]. Through long-term evolutionary processes, it has gained a vast number of glycosyl hydrolases, carbohydrate-binding proteins, and transporters, enabling it to utilize diverse carbohydrates [1]. Organized in discrete gene clusters, so-called polysaccharide utilization loci (PULs), *B. thetaiotaomicron* recognizes [2], binds to, degrades, imports, and metabolizes polysaccharides [3]. The glycolytic potential of *B. thetaiotaomicron* outnumbers most other human gut bacteria, with 92 predicted and verified PULs in its genome [4–18] (Suppl. File 1 PUL table). One of the best characterized gene clusters is the starch utilization system (SusRABCDEFG), enabling *B. thetaiotaomicron* to degrade starch, a glucan consisting of linear α-(1,4) linked amylose and the branched amylopectin with an α-(1,4) linked backbone and α-(1,6) branch points. A TonB- dependent transporter (TBDT), SusC [19], mediates the import of the substrate across the outer membrane. SusD, a surface lipoprotein is responsible for starch binding [20] along with SusEF [21]. SusG, an alpha-amylase, binds to starch and cleaves it into smaller oligosaccharides [22]. SusA and SusB, a neopullulanase and an α-glucosidase, respectively, further degrade these oligosaccharides in the periplasm into di- and monosaccharides [2]. Finally, SusR, a transcriptional regulator, modifies sus expression depending on the availability of starch or starch components [23]. Besides the classical SusC/D homologues in PULs other TBDT are predicted [24], which mediate, besides glycan uptake, vitamin and iron uptake [25, 26], summing up to 121 predicted TBDT overall. Additional important PUL proteins for *B. thetaiotaomicron* to thrive and diversify in the gut include glycosyl hydrolases, carbohydrate-binding proteins, sulfatases, polysaccharide lyases, carbohydrate esterases as carbohydrate-active enzymes (CAZymes). PUL expression is regulated by two-component system sensor histidine kinases/response regulators and extracytoplasmic function (ECF)-type sigma factors/anti-sigma factors [1]. Finally, transposons, integrases, and glycosyl transferases, which enable the building of phase variable capsular polysaccharides and TBDT, shape the genetic makeup and phenotype, respectively [1, 24, 27].

The metabolic diversity and adaptability to the environment of the gut and nutrient availability define the bacterial niche space and fitness within the gut community [28]. Thus, the ecological opportunity, e.g. changes in nutrient availability, can drive adaptive diversification. For example, Bacteroides fragilis diversifies through adaptive mutations in individual healthy human guts within complex microbiomes in the long-term [29]. Parallel mutations in cell envelope biosynthesis and polysaccharide utilization genes within one or multiple subjects arising from one unique ancestor strain each revealed that such pathways are under selection in a natural environment, suggesting novel selective pressures. Genetic changes in regulatory proteins can as well contribute to diversification and adaption to differential resource use, as exemplified by Escherichia coli. A transposon-mediated activation of a gene involved in acetate metabolism keeps this pathway active even when glucose is present, whereas the ancestral strain only slowly switches to acetate metabolism once glucose is depleted [30].

To evaluate the importance of nutrient adaptation in *B. thetaiotaomicron* and its specialization in complex carbohydrates, we exposed *B. thetaiotaomicron* to different nutrient landscapes with increasing complexity for three months: a simple medium with glucose in minimal medium (M1 medium), a medium with glucose and amylopectin (M2 medium), and a more complex medium with glucose, amylopectin, arabinogalactan, inulin, pectin, and xylooligosaccharides (XOS; M3 medium). We then evaluated whether adaptive diversification would occur in an ecologically meaningful time scale. Thus, we sequenced the evolved populations to identify mutational patterns. We observed that more complex media were able to support a higher number of mutations. We observed high levels of phase variation and confirmed gene clusters affected by site-specific integrase activity in SusC-like protein loci [24, 27]. We expanded those to N-acylglucosamine 2-epimerase and glutaminase A proteins. We found homologous recombination between type I restriction enzyme specificity protein genes leading to phase variation, as also recently suggested (PREPRINT, doi.org/10.1101/2024.02.17.580834). To verify changes in fitness, we competed the evolved populations and lineages isolated from evolved populations against the ancestor. Evolved populations outcompeted their ancestor, indicating rapid adaptive diversification. Glucose depletion rates in M3-evolved lineages were faster compared to the ancestor. During growth in the presence of both glucose and amylopectin, *susC^starch^* and *susD^starch^* were transcriptionally upregulated in M3- evolved lineages in comparison to the ancestor, indicating that evolved lineages de-regulate nutrient prioritization. In summary, *B. thetaiotaomicron* can rapidly respond to different nutrient environments through adaptive diversification that affects the regulation of nutrient utilization.

## Materials and Methods

### Bacterial strains and culture conditions

Bacteroides thetaiotaomicron type strain VPI-5482 was bought from DSMZ (DSM 2079). *B. thetaiotaomicron* isogenic type strain, evolved populations and lineages were either grown in brain heart infusion medium with supplements (BHIs, 37 g/l BHI, 5 g/l yeast extract, 1 g/l NaHCO_3_, 1 g/l L-cysteine, 1 mg/l vitamin K1, 5 mg/l hemin; details see Suppl. Table 3), or defined Minimal Medium (MM; see Suppl. Table 1) supplemented with 2.5 g/l glucose (Medium 1, M1), with 1.25 g/l glucose and 1.25 g/l amylopectin (Medium 2, M2) or with a carbohydrate mix (0.42 g/l glucose, 0.42 g/l amylopectin, 0.40 g/l arabinogalactan, 0.42 g/l inulin, 0.43 g/l pectin, and 0.42 g/l xylooligosaccharides; Medium 3, M3; for detailed information see Suppl. Table 2). For agar plates, 15 g/l agar was added. All populations and lineages were grown and competed at 37° C in an anaerobic chamber (Coy Labs, USA), under anaerobic conditions (85 % N_2_, 10 % CO_2_, 5 % H_2_).

Frozen stocks of the ancestor strains and the evolved populations and lineages were stored in 20 % glycerol, 80 % medium at −80° C, whereas the ancestor was stored in BHIs and the evolved populations and lineages in the respective medium they were evolved in.

### Generation of a tetracycline-resistant *B. thetaiotaomicron* strain

A derivative version of the *B. thetaiotaomicron* strain VPI-5482, ancestor^tetR^, carrying tetracycline resistance was constructed by inserting the Bacteroides species integrative vector pNBU2-bla-tetQb (Tet^R^, Amp^R^; kindly provided by Eric Martens, University of Michigan Medical School, USA) into one of two tRNA^ser^ attachment sites targeted by NBU2 [5, 31] (Suppl. Fig. 6a). This vector integrates DNA into the *B. thetaiotaomicron* genome in single-copy, without disrupting any genetic functions. E. coli S17-1 λ pir [32] was used as a donor for conjugation of DNA into *B. thetaiotaomicron*. For conjugation purposes, *B. thetaiotaomicron* was grown in BHIs medium and E. coli S17-1 λ pir was grown in LB medium supplemented with ampicillin (100 µg ml^-1^). Transconjugants were obtained by filter mating [33] and were selected in BHIs agar plates as tetracycline-resistant (Tet^R^, 3 μg ml^-1^ tetracycline) isolates. Growth of isolates was confirmed in BHIs medium + 3 μg ml^-1^ tetracycline (Suppl. Fig. 6b). Proper insertion of the vector into tRNA^ser^ locus 1 was verified by PCR using primer pairs listed in Suppl. Table 4 (Suppl. Fig. 6c). 

### Experimental evolution experiment

An isogenic culture of *B. thetaiotaomicron* (VPI-5482), referred to as ancestor, was grown in MM with 0.5 % (w/v) glucose. The isogenic *B. thetaiotaomicron* strain was subcultured into MM supplemented with different carbohydrate sources (M1, M2, or M3, media composition see above) with 7 replicates each. Serial transfers of *B. thetaiotaomicron* were carried out daily for forty-eight days, by transferring 100 µl into 5 ml new media. A negative control for each medium was included. All experiments were carried out under anaerobic conditions. Glycerol stocks of each evolved population were made once a week.

### Isolation of evolved lineages

100 µl bacterial culture from a glycerol stock of M1-M3-evolved populations (n=4 replicates each medium) were inoculated into 5 ml medium, in which the populations were evolved. Bacteria were grown o.n. at 37° C. Serial dilutions in PBS of the bacterial cultures were carried out and 100 µl of a 10^-5^ dilution were plated on M1-M3 agar plates. Colonies were grown for three days. To ensure clonality, ten randomly picked colonies were restreaked on new agar plates and grown for an additional two days. M1-M3-evolved colonies were inoculated in 5 ml M1-M3, respectively and incubated o.n. at 37° C. Glycerol stocks of the o.n. cultures from the lineages were frozen at −80° C.

### DNA isolation and library preparation for shotgun metagenome sequencing of evolved populations

We performed whole genome shotgun sequencing for the ancestor and 7 replicates of M1-, M2-, and M3-evolved populations. Evolved populations were grown to an optical density OD_600 nm_=0.5-1 in the medium they were evolved in. One ml of the bacterial culture was pelleted at 15,000 g for 3 min at 4° C. DNA was isolated using the All-DNA/RNA/Protein prep Kit for bacteria (Qiagen), according to the manufacturer’s instructions. DNA concentration was determined using Quant-iT PicoGreen dsDNA Assay Kit (Thermo Fisher Scientific) according to the manufacturer’s protocol. Fluorescence was measured on a Tecan reader using 480 nm excitation and 520 nm emission wavelength.

For library preparation we followed the TruSeq DNA PCR-free Kit (Illumina) according the manufacturer’s instructions for 350 bp insert size. DNA shearing was performed on a Covaris S220 Series using 100 ng DNA input in a volume of 130 µl. DNA concentration was determined using Qubit dsDNA HS Assay Kit (Thermo Fisher Scientific). Metagenome sequencing was performed on a NextSeq550 System (Illumina) with 150 bp paired-end reads at the VBCF (Vienna BioCenter Core Facilities).

### Lineage sequencing

M3-evolved lineages were grown in M3 medium o.n.. One ml of the o.n. culture was pelleted at 8,000 g and washed once with PBS. DNA was isolated using the Wizard Genomic DNA Purification Kit (Promega), according to the manufacturer’s instructions for isolation of Gram negative bacteria. DNA concentration was determined fluorometrically on the Qubit 4.0 fluorometer using the Qubit dsDNA HS Assay Kit (Thermo Fisher Scientific). DNA fragment size was analyzed on a genomic DNA Tape (Agilent) on a TapeStation 4100. Barcoded genomic libraries were prepared using the NEBNext® Ultra™ II FS DNA Library Prep Kit for Illumina and sequenced on an Illumina MiSeq using MiSeq reagent kit v3 (600 cycles, 2x 300 bp mode).

### Sequence data preprocessing and mutational analysis

We used fastp for adaptor trimming and QC filtering of sequencing reads, with the following parameters: for NextSeq reads: -q 20 --detect_adapter_for_pe -l 80 --cut_tail --cut_tail_window_size 1; for MiSeq reads: -q 20 --detect_adapter_for_pe -l 80 --cut_tail --cut_tail_window_size 1 -- trim_tail2 40; 

Polymorphisms calls were made with the breseq v38.1 pipeline [34] in polymorphism mode for population samples. All calls were made against the RefSeq genome for *B. thetaiotaomicron* VPI-5482 with accession number; NC_004663.1 and plasmid with accession number; NC_004703.1. Polymorphisms had to be supported by at least 2 forward and 2 reverse reads and be detected in at least 0.01 frequency in the population. Additionally, we did not consider any fixed polymorphisms (>= 0.9 frequency) in the ancestral population. We masked any polymorphisms involved in phase variable regions, to exclude potential false positive calls they might have introduced. We used all available annotation data from NCBI’s RefSeq database. All RefSeq genome and annotations files can be accessed here:

https://ftp.ncbi.nlm.nih.gov/genomes/refseq/bacteria/Bacteroides_thetaiotaomicron/all_assembly_versions/GCF_000011065.1_ASM1106v1/

### Genome assembly of lineages

Individual read library bam files were merged using samtools v1.8 and were quality checked using fastQC v0.12.1. Adapters were trimmed and phiX contamination was removed using BBDuk (part of BBMap v39.01) to a minimum of 50 base pairs in length. Reads were k-trimmed from the right with a kmer of 21, minimum kmer of 11 and hamming distance of two along with the “tpe” and “tbo” options. Quality trimming was performed from the right with a Q-score of 15. BBMap’s reformat.sh was used to interleave read libraries and libraries were merged. The interleaved read library was assembled using SPAdes v3.15.1 in “isolate” mode with kmers set to 21, 31, 41, 51, 61, 71, 81, 91, 101, 111, 121. BBMap’s reformat.sh was used to remove contigs under 1000 base pairs from the assembly. The assemblies were binned into putative genomes using MetaBAT2 v2.15. Coverage information was calculated using BBMap v39.01 with a 98 % identity and the resulting bam file was sorted using samtools v1.16.1. Metabat2 was run with a minimum length of 1500 base pairs and default workflow was used in concoct. Only the sample’s readset was used to provide coverage information. Putative genomes were also binned using Metabat2 with no coverage information. A quality comparison of the original assembly and all putative MAGs was performed using checkM v1.2.2 lineage workflow alongside coverage information from BBMap (at 95 % identity). If the assembly was determined to be low contamination (<10 %), it was used in later analysis. If the assembly was too contaminated, a MAG of high quality and high coverage (representing the dominant isolate in the sample) was chosen for later analysis. The assemblies and MAGs chosen were annotated using prokka v1.14.6.

### Shufflon identification

We selected regions with polymorphisms with highly correlated frequency trajectories, in a relative small genomic distance (100 bp), as signatures for phase variation. We used genome assemblies of the lineages to verify these regions. We included assemblies which passed our quality criterion of N50 > 100K. Genes were identified with Prodigal v2.6.3 [35] and were associated to the reference genome according to their best blastn hits. Concordance fraction between genes orientation on the contig and the reference genome, determined contig’s orientation. When concordance was < 50 %, contig orientation was reversed. To call a phase variable region (or shufflon), at least one recombination event should be detected, after including only assemblies where all genes involved would be physically linked (present on the same contig).

### Shufflon verification in RefSeq *B. thetaiotaomicron* genomes

All nineteen complete *B. thetaiotaomicron* genomes available in the RefSeq database as of January 2024 were used to identify shufflon signatures. Construction of locally collinear blocks was performed with a 500 nt threshold of minimal block length using SibeliaZ [36] and Maf2Synteny [37] integrated into the Badlon pipeline. The Panacota modules were run with the following parameters: at step prepare only complete genomes were included; step annotate was performed using Prokka v1.14.6 [38] and Prodigal v2.6.3 [35]; step pangenome was performed with 60 % protein similarity threshold; step corepers was used to select single-copy universal genes; step align was performed using MAFFT v7.520 [39]; step tree inferred a phylogenetic tree from the concatenation of alignments of single-copy universal genes using IQTree version 2.2.0.3 [40] with 1000 bootstraps and the GTR DNA substitution model. The tree was visualized with iTOL version 6.7.4 [41]. Then the structural variants were reconstructed using PaReBrick v.0.5.5 [42].

### *p_N_p_S_* calculations

We calculated *p_N_p_S_* directly from NGS data with a Python script using previously published methodology [43]. To reach the quality, we filtered the alignment out for reads with Quality Control (QC) ≥ 30 and Mapping Quality (MQ) ≥ 20, while discarding any duplicates or secondary reads. Finally, only codons supported completely by at least four reads were used in the calculation.

As the number of observed substitutions in the experiment was not enough for accurate p_N_p_S_ calculation for individual genes (mean value ∼2.3 per ORF), we used concatenated sequences to compare cohorts of genes, e.g. the ones within and outside PULs.

### Competition assay of evolved populations, evolved lineages and ancestor

In order to perform competition experiments *B. thetaiotaomicron_(ancestor)_* was genetically modified to include a tetracycline gene (*B. thetaiotaomicron^ancestor-tetR^*, ancestor^tetR^ Suppl. Fig. 6). If not stated otherwise, evolved populations were pre-grown in the respective medium they were evolved in, the ancestor or the ancestor^tetR^ in BHIs. Bacteria were grown under anaerobic conditions for 6-7 h to reach exponential growth phase. Bacteria were diluted to an optical density OD_600 nm_ of 0.10-0.12 in the medium they will be competed in and 300 µl of each evolved population or lineage was competed against the ancestor^tetR^ in a 1:1 ratio in 5 ml of fresh medium for 15 h-17 h or 24 h. The ancestor and the evolved population were competed in either BHIs, M1, M2, M3, or MM with a single carbohydrate of the complex M3 medium. One ml of each bacterial culture was harvested at competition start (0h) and at experiment end in stationary phase for DNA extraction.

### DNA extraction and quantitative PCR for competition experiments

One ml of bacterial culture was centrifuged at 10,000 g for 10 min and washed in 1 ml PBS. The bacterial pellet was resuspended in 100 µl nuclease-free water and heated for 10 min at 95° C with repeated vortexing. The cell suspension was centrifuged at 13,000 g for 3 min and the DNA-containing supernatant was frozen at -20° C until further analysis.

Quantitative PCR was carried out for the housekeeping gene RNA polymerase beta (rpoB), present in all bacteria, and the tetracycline resistance gene TetQ, which is only present in the genetically engineered ancestral *B. thetaiotaomicron* strain (ancestor^tetR^). The DNA was diluted 1 by 20 and 1 μl of the dilution was subjected to quantitative PCR using 0.4 μM of primers for TetQ or rpoB (primer sequences can be found in Suppl. Table 5) and 1x iQ SYBR Green Mix (BioRad) in a total reaction volume of 20 μl. qPCR was performed on a CFX96 Real-Time PCR Detection System (BioRad) with following cycling conditions: 95° C for 5 min, followed by 40 cycles of 95° C for 15 s, 56° C for 20 s, and 72° C for 30 s. To determine the specificity of PCR reactions, a melting curve analysis was carried out after amplification by slowly cooling from 95° C to 60° C, with fluorescence collection at 3° C intervals and a hold of 10 s at each decrement.

### Calculation of fitness advantage of the evolved population and lineages against the ancestor

To calculate fitness advantage (FA) resulting from the competition, an adaption of the ΔΔCt method was used

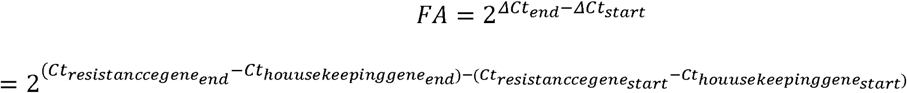

whereas Ct is the threshold cycle for gene detection upon qPCR, a change in Ct of one is equal to a doubling of bacteria, which is equal to a generation. *ΔCt_end_* is the difference in Ct value of the tetracycline resistance gene, which is only present in the ancestor^tetR^, and the Ct value of the housekeeping gene, which is present in all bacteria, at the end of competition. *ΔCt*_start_ is the difference in the Ct value of the tetracycline resistance gene and the Ct value of the housekeeping gene, directly after inoculation and start of the competition. Data was removed from analysis if Ct values were >30, which indicates low inoculation concentration of bacteria and if ΔCt_rpoB_=Ct_start rpoB_-Ct_end rpoB_ values were <3, which indicates overall bad growth. To be able to quantify proper FA values, a minimum growth threshold for the ancestor^tetR^ was determined and set to ΔCt_tetR_=Ct_start tetR_-Ct_end tetR_<1. Below that threshold, it is an all-out win situation for the competitor, meaning, the competitor (evolved population or lineage) outgrew the ancestor^tetR^. All-out wins by the evolved strains were set to a FA value of 100.

If the FA value is around 1, no FA is present. Pre-growth conditions in different media slightly altered competition performance and thus FA by a factor of 2 (p=0.036; Suppl. Fig. 7). Hence, we used a threshold of 2 to incorporate this variation to distinguish populations with a competition advantage from those without (for details see supplementary results). For further experiments, the ancestor^tetR^ was pregrown in BHIs.

### RNA isolation, cDNA synthesis and quantitative real-time PCR

RNA was extracted using the Total RNA Purification Kit (Norgen Biotek). DNase treatment was performed using Turbo DNase (Thermo Fisher Scientific) and RNA was purified using the Zymo RNA Clean & Concentrator Kit (Zymo Research). RNA was quantified with the Qubit 4 fluorometer and Qubit RNA BR reagents (Thermo Fisher Scientific). cDNA was synthesized according to the High-Capacity cDNA Reverse Transcription Kit (Thermo Fisher Scientific). qPCR was performed as stated above using primers for SusC, SusD and the control gene 16S (primer sequences can be found in Suppl. Table 5). The ΔΔCt-method was used to calculate relative expression.

### Glucose depletion

One ml of bacterial culture was spun at 10,000 g for 5 min. The supernatant was transferred into a new tube and potentially remaining cells were killed at 65° C for 10 min. For glucose measurement of the culture supernatants, the ACCU-CHEK guide (Roche) was used. 10 µl of the supernatant were transferred onto an ethanol-cleaned glass slide and the glucose test strip was put into the liquid until measurement occurred.

### Cell size determination

The ancestor, ancestor^tetR^ or M1-M3-evolved lineages were pregrown for 6-7 h in BHIs or in the medium they were evolved in, respectively. Bacterial cells were diluted to an OD_600 nm_=0.1 in M3 medium, and 300 µl diluted bacteria were used for competition in 5 ml M3 medium, or 600 µl if grown alone. Cells were grown for indicated times. At each point in time 10 µl of bacterial culture were transferred onto a microscope slide with 6 mm reaction wells (MARI1216690, Marienfeld), air- dried, fixed in 2 % paraformaldehyde o.n., washed in PBS and bacterial DNA was stained with 10 µl 1 µg/ml DAPI in PBS. Slides were dipped in ice-cold H_2_0 and air-dried. Slides were embedded in Citifluor and images were taken on a CLSM (Leica TCS SP8X) using the Leica Application Suite X. DAPI images were recorded with a 405 nm laser diode, a HC PL APO CS2 93x/1.30 glycerol objective, a HyD detector, a pinhole size of 1 AU, a resolution of 2328x2328 pixels and a size of 98.04 x 98.04 µm. For image deconvolution, Lightning was applied. Bacterial cells were measured using ImageJ2/Fiji (NIH). The Feret’s diameter in µm (the longest distance between any two points along the selection boundary of a cell, also known as maximum caliper) was determined using the analyze particle function. Particles were manually screened to select single cells only, cells on the image edges were excluded from analysis.

### Statistics

Statistical analysis was performed using R statistical software (https://www.r-project.org/). For pairwise comparisons of FA, Feret’s diameter of bacterial cells, and OD values Kruskal-Wallis test and for multiple comparison correction the false discovery rate (FDR) method was used. Kruskal-Wallis rank sum test with Dunn’s correction was used to reveal significant changes in number of mutations/nucleotide between M1-M3 evolved populations for any PUL gene or all genes. Significant differences in number of mutations/nucleotide between PUL gene or all genes, independent of the media the population replicates have evolved in, were analyzed using Aligned Ranks Transformation ANOVA with Tukey’s post-hoc correction. Statistical significant differences between percentage of compromised cells during competition and without competition were calculated using Independent Samples T-test. Paired Wilcoxon signed-rank test was used to investigate differences in pregrowth conditions (BHIs and M3 medium) of ancestor^tetR^ and evolved populations or ancestor during competition experiments. To compare variances of evolved populations against variances of evolved lineages F-test was used. Repeated measures ANOVA with Bonferroni correction was performed to compare FA of M3-evolved populations in M3 medium versus single carbohydrates thereof. To calculate differences in susC/D expression the area under the curve was calculated, followed by an ANOVA with Tukey’s HSD correction. To calculate differences in glucose depletion between the ancestor and M3-evolved lineages Friedman rank sum test using Conover’s single-step correction was used. Adjusted p-values were considered statistically significant if less than 0.05 (*p⍰<⍰0.05; **p⍰<⍰0.01; ***p⍰<⍰0.001). For some graphs a lettering system was used to denote statistically significant data, whereas different letters indicate statistical significant differences between groups.

## Results

### Site-specific integrase-mediated gene shuffling in evolved populations

To better understand how commensal gut bacteria adapt to different carbohydrate landscapes, we studied the genetic diversification of *B. thetaiotaomicron* in defined nutrient conditions with varying carbohydrate substrate complexity. An isogenic population of *B. thetaiotaomicron* was serially passaged daily for 84 days in a defined minimal medium (MM, Suppl. Table 1) supplemented with (1) glucose (M1), (2) glucose and amylopectin (M2), or (3) six different carbohydrates (glucose, amylopectin, arabinogalactan, inulin, pectin, and xylooligosaccharides [XOS]; M3; Fig. 1, Suppl. Table 2). Cultivation conditions allowed for approximately six doublings a day, as determined by calculating generation numbers using qPCR, resulting in approximately 500 generations during the entire experiment.

**Fig. 1.**
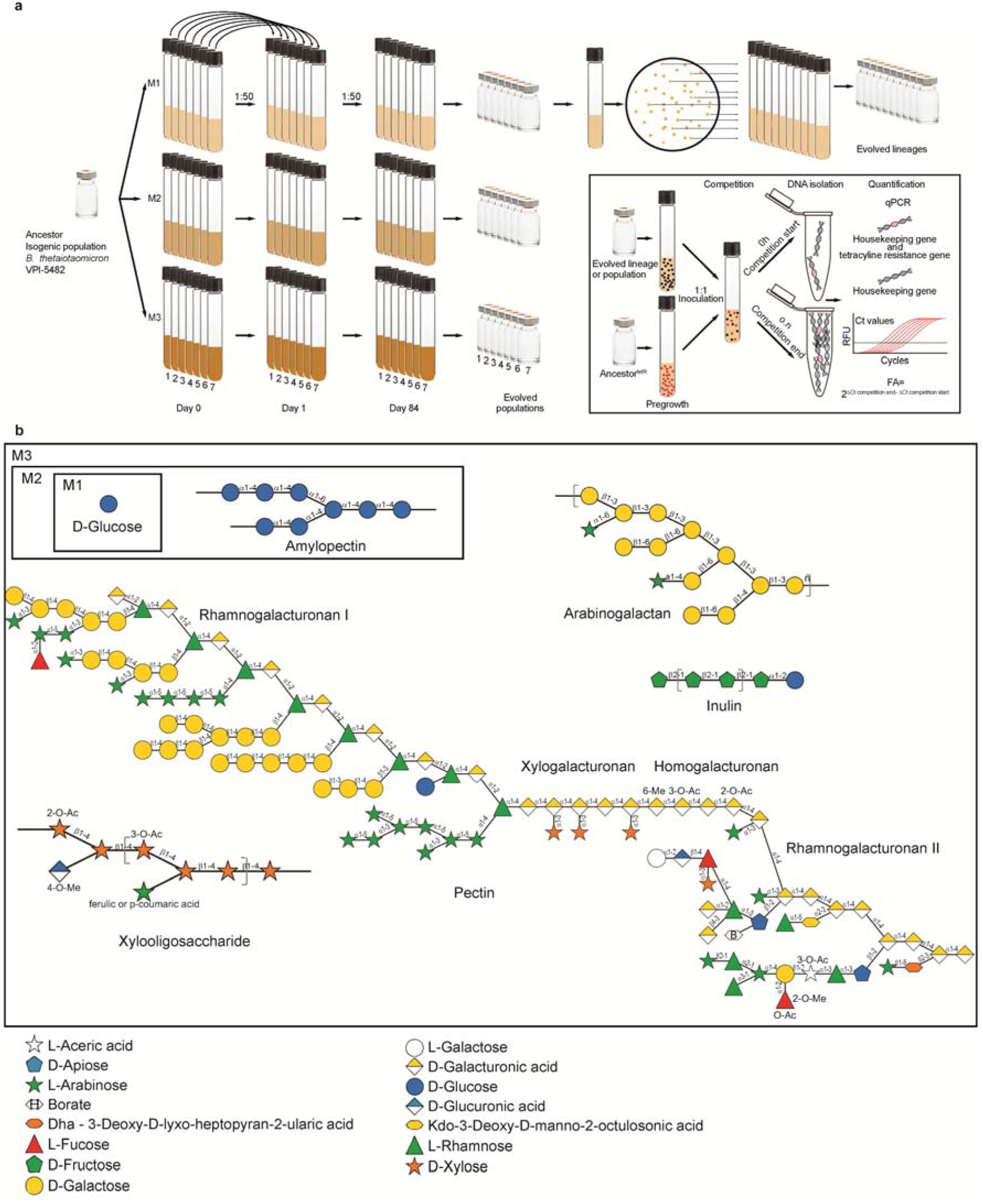
Experimental setup and glycan structures. **a** An isogenic *B. thetaiotaomicron* strain VPI-5482 (ancestor) was serially diluted for forty-eight days in 7 replicates of Medium 1 (MM containing glucose, M1), Medium 2 (MM containing glucose and amylopectin, M2), or Medium 3 (MM containing glucose, amylopectin, arabinogalactan, inulin, pectin, and XOS, M3). Glycerol stocks of evolved populations were made and stored at −80° C. Evolved populations from glycerol stocks were cultured on the respective media agar plates they were evolved in, restreaked to insure clonality, and frozen stocks were made from evolved lineages. Evolved populations and lineages were used for competition experiments against an ancestor *B. thetaiotaomicron* strain with a tetracycline resistance gene incorporated in its genome (ancestor^tetR^). After a pre-growth phase of 5-6 h in the media bacteria were evolved in and after which bacteria are in logarithmic growth phase, bacterial populations or lineages were inoculated 1:1 with the ancestor^tetR^, which was pre-grown in BHIs. At the start of competition and after o.n. incubation in stationary phase bacterial suspensions were collected, DNA was isolated, and subjected to quantitative PCR. qPCR was performed for a housekeeping gene, present in all bacteria (black dots and DNA strands) and for the tetracycline resistance gene, present in the ancestor^tetR^ only (red dots and DNA). Fitness advantage (FA) levels of the evolved populations and lineages against the ancestor were calculated using Ct values as mentioned in the Material and Methods section. **b** Glycans used in this study were drawn with DrawGlycan-SNFG (Symbol Nomenclature for Glycans) [44]. D-Galacturonic acid in pectin can be modified by methyl (6-Me) and acetyl (2-O-Ac, 3-O-Ac) esterification [45]. XOS can be further substituted at the 2’-OH or 3’-OH position with acetyl or 4-O-methyl glucuronyl groups or arabinose [46–48]. Arabinose itself can be esterified with ferulic - or p-coumaric acid.

In the initial screening for mutations in the evolved populations, we identified haplotype-like signatures of single nucleotide polymorphisms (SNPs) that could be introduced due to phase variation in these regions (Zioutis et al, in preparation). After inspection of genome assemblies from clonal lineages, we confirmed the presence of one experimentally verified and one proposed shufflon involving genes encoding SusC-like proteins BT1040-BT1042-BT1046 [27] and BT2260-BT2262- BT2268 [24], respectively (genomic localization of shufflons is outlined in Fig. 2).

**Fig. 2.**
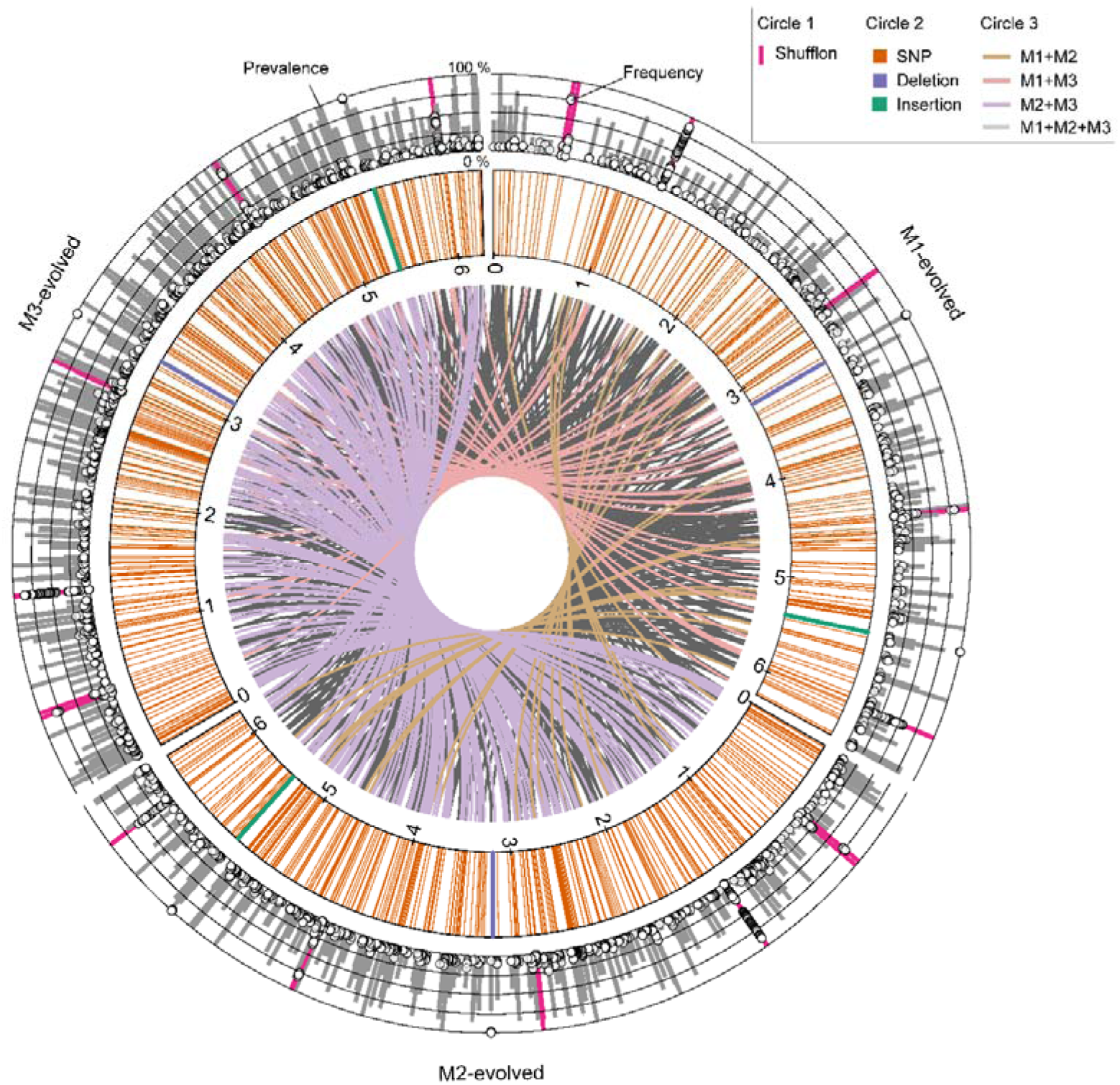
Circos plot of genomic variation in M1-M3 evolved populations. Circos plot illustrating SNPs, insertions, and deletions on the *B. thetaiotaomicron* genome of M1-M3-evolved populations as well as medium prevalence of the mutation (grey bars) and frequency of the mutation in every population replicate (circles) in circle 1. Only polymorphisms, which are present at least at 5 % frequency and found in at least three replicates are presented in the circles. Shufflon regions are denoted as pink bars. Circle 2 shows the distribution of SNPs (orange), insertions (green) and deletions (purple) within the genome. The connecting lines in the innermost circle (Circle 3) point out mutations which are found in populations of at least two media, which are defined by different colours, according to the media.

In addition, we identified two other recombinational shufflons. One involved PUL8 and PUL9 (Suppl. File 1 PUL table), leading to gene order and orientation changes due to the inversion introduced at N- acylglucosamine 2-epimerase genes (BT0437, BT0453), but did not affect the coding sequence (Fig. 2). The other shufflon involved glutaminase A genes BT3477, BT3481, BT3491, and BT3503 (Fig. 2). Furthermore, we identified sites of recombination involving type I restriction modification enzymes, specifically the DNA specificity subunits BT4540, BT4541, BT4542, and BT4543 (hsdS gene, Pfam domain PF01420) (Fig. 2), which has recently been suggested to be phase-variable (PREPRINT, doi.org/10.1101/2024.02.17.580834). Analysis of all currently available complete *B. thetaiotaomicron* assemblies in RefSeq supports the presence of phase variation in four out of five of the identified loci in this study. These loci have also been observed to be phase variable during evolution of *B. thetaiotaomicron* in mono-colonized mice in a parallel study conducted by our group (Zioutis et al, in preparation), indicating high rates of gene rearrangements both *in vitro* and *in vivo*. The identification of these gene rearrangements in very different conditions may indicate that they are high-frequency variable regions that are not necessarily under selection pressure in the current experiment.

### Conserved and medium-specific genetic diversification

In addition to the structural variants described above, most genetic variants identified were single nucleotide polymorphisms (SNP, 99.3 %), followed by deletions (DEL, 0.5 %) and insertions (INS, 0.2 %) (Fig. 2). Intergenic mutations accounted for approximately one-sixth of all mutations. From a total of 3,397 mutations, 435 (12.8 %) were identified in at least one replicate of all three media types (Fig. 3a). 1,367 (40.2 %) mutations were unique to M3-evolved populations, 573 (16.9 %) were unique to M2-evolved populations, and 224 (6.6 %) to M1-evolved populations. Of note, we found that the number of mutations increased with the complexity of the medium (Fig. 3c). Shared mutations were found mostly between populations evolved in M2 and M3 (564, 16.6 %), although this may in part be influenced by the variable number of replicates analyzed, as three sequencing libraries (2 M1 and 1 M3 replicate) produced insufficient read depth and were excluded from further analysis (Fig. 2 and 3a).

**Fig. 3.**
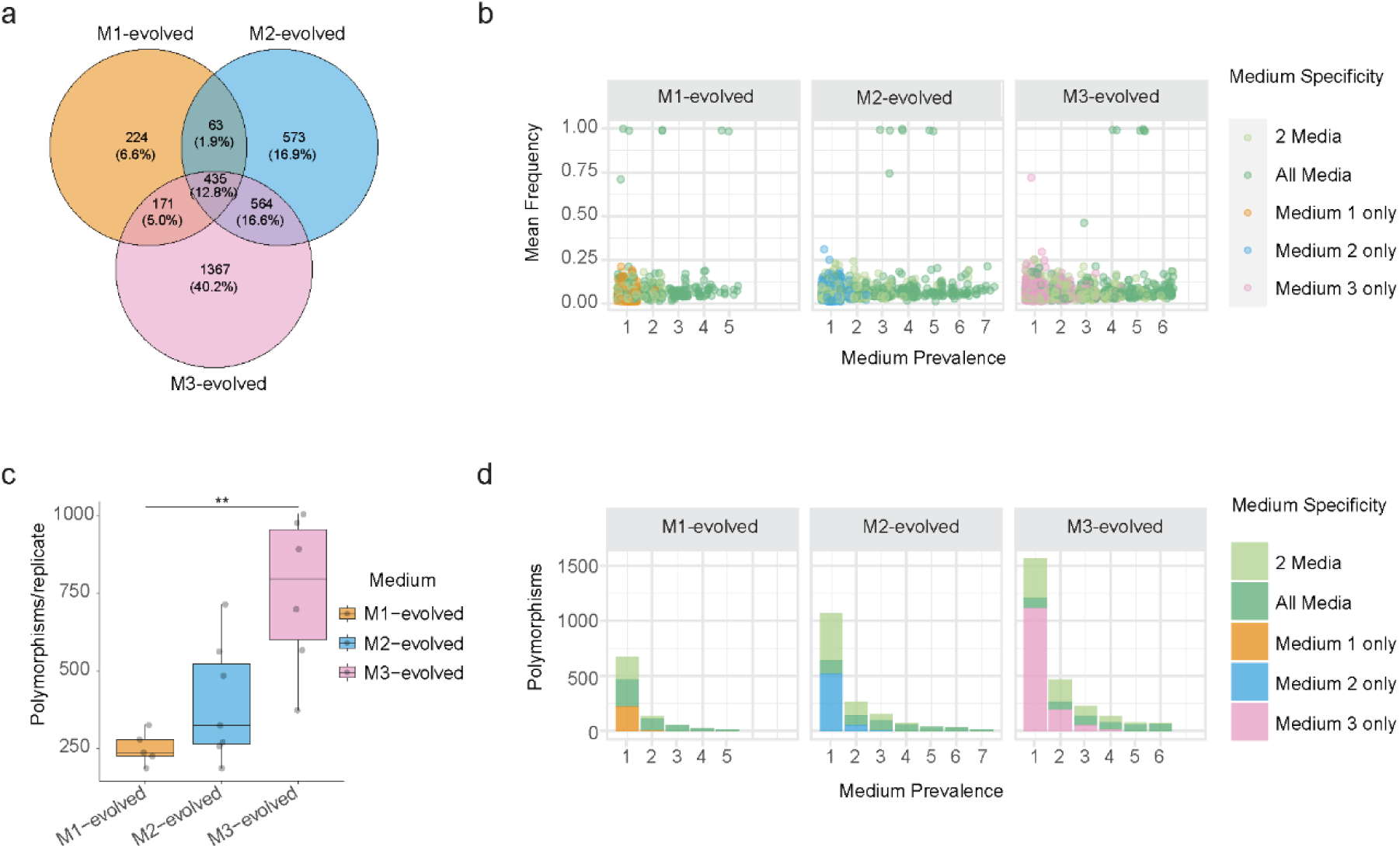
Polymorphisms of M1-M3 evolved populations. **a** Venn diagram representing polymorphisms which are unique for one medium-evolved population or common in two or three different media-evolved populations. **b** Mean frequency of polymorphisms only present in a specific medium (M1: orange, M2: blue and M3: pink dots) or more than one medium (green dots) for the three media-evolved populations and the prevalence, i.e. the number of replicates each media in which they are present. **c** Number of polymorphisms each replicate (grey dots) for the different media evolved populations (n_M1-evolved_=5, n_M2-evolved_=7, n_M3-evolved_=6). Significant differences of the number of polymorphisms between two groups were analyzed using Kruskal-Wallis test, Bonferroni corrected (**p-value<0.01). **d** Total number of polymorphisms for all replicates of the three media-evolved populations in a specific medium (M1: orange, M2: blue and M3: pink) or more than one medium (green) and the medium prevalence.

Six mutations segregated in frequencies approximating fixation in the population. They were intergenic and present in evolved populations from all three media, although not always present in all replicates (Fig. 3b). 99.1% of mutations were low to moderate frequency, with a mean frequency below 25%. A substantial fraction (36.3%) of polymorphisms were shared between two or three different media-evolved populations (Fig. 3a, d), with varying degree of medium prevalence, i.e. the presence in a replicate of one specific medium-evolved population. The parallel appearance of the same polymorphism across replicate populations has been suggested to be indicative of adaptive evolution [49]. We therefore next evaluated the parallelism of polymorphisms in the three media conditions. We found that medium-specific polymorphisms were detected in evolved populations from all three media. Among these, polymorphisms were more likely to be detected in multiple replicates for populations evolved in more complex media (Fig. 3d). Polymorphisms that were unique to a specific medium only appeared in up to two out of five (40 %) M1-evolved populations, up to three out of seven (43 %) M2-evolved populations, and up to four out of six (67 %) M3-evolved populations.

Out of 92 experimentally proven and putative PULs (Suppl. File 1 PUL table), 79 showed at least one mutation in any gene in the PUL in at least one evolved population. The remaining 13 PULs with no mutations have either unknown substrates or degrade mucin O-glycans [5], which were not present in the media. On average, 17 % of all polymorphisms detected were in PULs. As PULs themselves make up approximately 19.3 % of the protein-coding genome [50], no increased occurrence of mutations within PULs was observed. However, when calculating the ratio of nonsynonymous versus synonymous substitutions, PUL genes presented with an elevated dN/dS ratio compared to all other *B. thetaiotaomicron* genes (in all biological replicates) indicating positive selection of PUL genes independent of the media [51] (Suppl. File 2). A closer evaluation of polymorphisms found within PULs revealed many mutations in carbohydrate esterases, polysaccharide lyases, sulfatases and hybrid two-component systems (HTCS), the latter being regulatory proteins that induce expression of the respective PUL upon carbohydrate availability (Suppl. Fig. 1a). However, only carbohydrate esterases showed a significantly enriched number of mutations compared to all genes (Suppl. Fig. 1a, b). In contrast, mutations were rare in ECF-type sigma factor/anti-sigma factor, the other main type of PUL regulatory system which are more often found in PULs important for host-derived-glycan degradation (Suppl. File 1 PUL table). SusC-like family genes and carbohydrate esterases were among the gene families most enriched in M2- and M3-evolved populations compared to M1-evolved populations. The carbohydrate esterases that accumulated polymorphisms were a xylan esterase (CE4, EC3.1.1.72), which removes acetyl groups from XOS (see Supplementary Results) as well as (sialate 9-O-) acetylesterases (CE20, EC3.1.1.53 and 3.1.1.6), which remove acetyl groups from N- acetyl- or N-glycoloyl-neuraminic acid and also cleave off other acetyl groups found in rhamnogalacturonan II and XOS [52]. SusD-like proteins, which mediate carbohydrate binding, accumulated fewer polymorphisms compared to all genes. Collectively, this data suggests mutational hotspots in PULs and potentially rapid adaptive diversification of *B. thetaiotaomicron* to different nutrient environments (for more examples see Supplementary Results).

### Evolved populations have increased fitness

We next performed competition experiments to determine if the genetic diversification of evolved populations confers increased fitness. During competition in M1 medium, populations evolved in all media showed a significant fitness advantage (FA, see Methods) compared to the ancestor (mean FA = 4.8-6.6; Fig. 4a). Evolved populations also strongly outcompeted the ancestor in M2 medium (mean FA=22-60; Fig. 4a). During competition in M3 medium, M2- and M3-evolved populations show the highest FA (mean FA=61 and 55, respectively; Fig. 4a). M1-evolved populations, however, had a similar FA as if competed in M1 medium. An increased efficiency of utilizing carbon sources in M3 medium is also reflected by the elevated optical density of M2- and M3-evolved populations at stationary phase (Suppl. Fig. 2a). These results indicate that M2- and M3-evolved populations have the best adaption to M3 medium. Of note, in a complex medium (BHIs), the FA of all M-evolved populations remained similar to the ancestor (Fig. 4a).

**Fig. 4.**
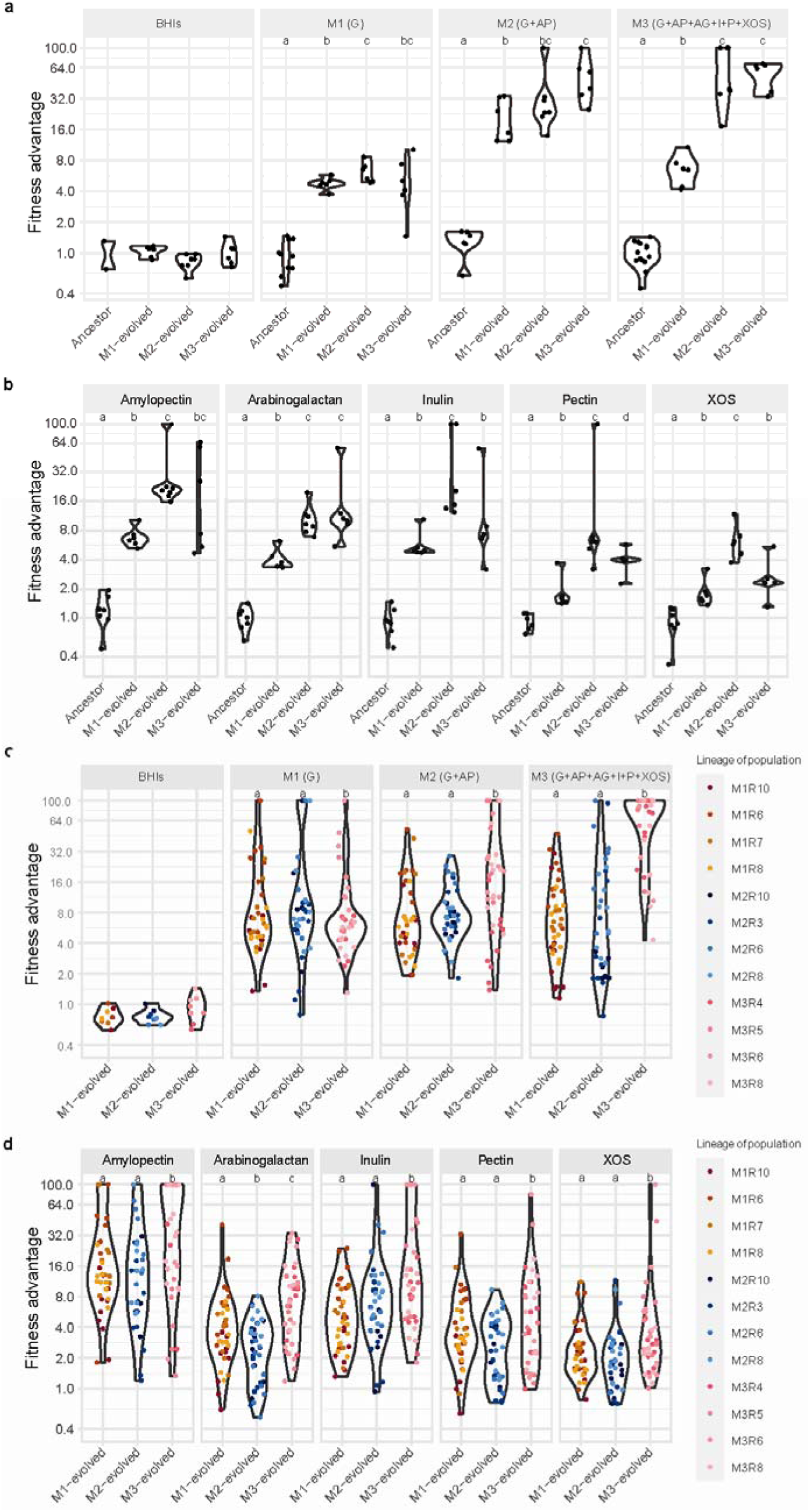
FA of evolved populations and lineages competed against the ancestor strain. Violin plot of the FA of the ancestor and M1-M3-evolved populations or lineages. Every dot represents the FA of the (**a, b**) ancestor or an evolved population and (**c, d**) evolved lineages competed against the ancestor^tetR^ in different media. Significant differences in FA between two groups were analyzed using Kruskal-Wallis test and are indicated by different letters. FA of M1-M3-evolved populations competed in (**a**) BHIs, M1-M3 medium (G, glucose; AP, amylopectin; AG, arabinogalactan; I, inulin; P, pectin; XOS, xylooligosaccharides), or (**b**) with single glycans of the complex M3 medium in MM. FA of M1-M3-evolved lineages competed in (**c**) BHIs, M1-M3 medium, or (**d**) with single glycans of the complex M3 medium in MM. Colours indicate different lineages from various replicates.

### FA of evolved populations is substrate-specific

We next evaluated the FA of evolved populations on the individual carbohydrates present in M3 medium. Competition in amylopectin resulted in high FA values (mean FA; M1=7, M2=31, M3=30; Fig. 4b), with M2-evolved populations showing higher FA values compared to M1 populations (p=0.003). Of note, the FA of M3-evolved populations was lower when competed in the presence of individual carbohydrates such as glucose, pectin, and XOS (p_adj_-value=0.022, p_adj_-value=0.012, p_adj_- value=0.01, respectively) but was not significantly different to amylopectin, arabinogalactan, and inulin, as sole carbon sources, in comparison to competition in M3 medium (Fig. 4b). For amylopectin, arabinogalactan, and inulin there was also elevated optical density of stationary phase cultures (Suppl. Fig. 2a, b). Although M2-evolved populations were not exposed to arabinogalactan, inulin, pectin and XOS their FA was not significantly different or higher from those of M3-evolved populations (p=0.39, p=0.01, p=0.02, p=0.009, respectively). Also, M1-evolved populations had an FA when competed on arabinogalactan and inulin compared to the ancestor (p=0.004, p=0.005, respectively). Overall, the lowest FA was seen for competition in pectin and XOS, with M1-evolved populations not showing any significant FA (Fig. 4b). Stationary phase optical density was also lowest on these carbohydrates, indicating low utilization efficiency (Suppl. Fig. 2b). In summary, competition experiments with single carbohydrates showed that FA in evolved populations was high when competed on amylopectin, arabinogalactan, and inulin. There was a trend for M2- and M3-evolved populations to have a higher FA compared to M1-evolved populations, although not always statistically significant.

### Evolved lineages have variable fitness

We next evaluated the fitness of clonal lineages isolated from genetically heterogeneous evolved populations (Fig. 1a). Eight to ten lineages isolated from four different evolved populations from each medium were competed against the ancestor in up to nine different media, resulting in a total of 964 competition experiments. Overall, most evolved lineages showed an FA when competed in M1, M2, or M3, but not in BHIs medium (Fig. 4c). In comparison to the evolved populations, the variation among FA of lineages showed sometimes higher and sometimes lower variance or equal variances (see Suppl. File 3 and 4 for p- and F-values). Competition in M1 resulted in a similar average FA between the M1-M2 evolved lineages, with a mean FA of 16 and 25, respectively. M3-evolved lineages showed a lower FA, with a 10-fold increase. During competition in M2 or M3, M3-evolved lineages had the highest FA, whereas no difference in FA for M1- and M2-evolved lineages was present. Also, M3-evolved lineages had higher optical density at the competition endpoint in M3 medium (Suppl. Fig. 2c).

Evolved lineages were also competed against the ancestor with single carbohydrates. As expected, and like the evolved populations, evolved lineages performed well in amylopectin (Fig. 4d). Overall, competition for inulin resulted in the second-highest FA values, especially for M3-evolved lineages. As expected, FA levels during competition in pectin and XOS were rather poor, with M3-evolved lineages reaching the highest levels of 9 and 10, respectively. Several lineages did not show a significant FA. In summary, evolved lineages have a similar mean fitness compared to evolved populations. Some lineages show no FA and others have higher FA than the evolved populations from which they were derived. The FA of some lineages during competition in certain carbohydrates was highly variable (Suppl. File 4), as was the stationary-phase optical density (Suppl. Fig. 2d).

Several evolved populations and lineages outcompeted the ancestor completely by inhibiting ancestor growth (Suppl. Fig. 3 and 5). The reason for the inhibition of the ancestor growth during competition is unclear. It might involve nutrient limitation as the drop in cell numbers corresponds to limited glucose availability (Suppl. Fig. 5d). Also, a direct effect of the evolved lineage on cellular integrity of the ancestor cannot be excluded.

**Fig. 5.**
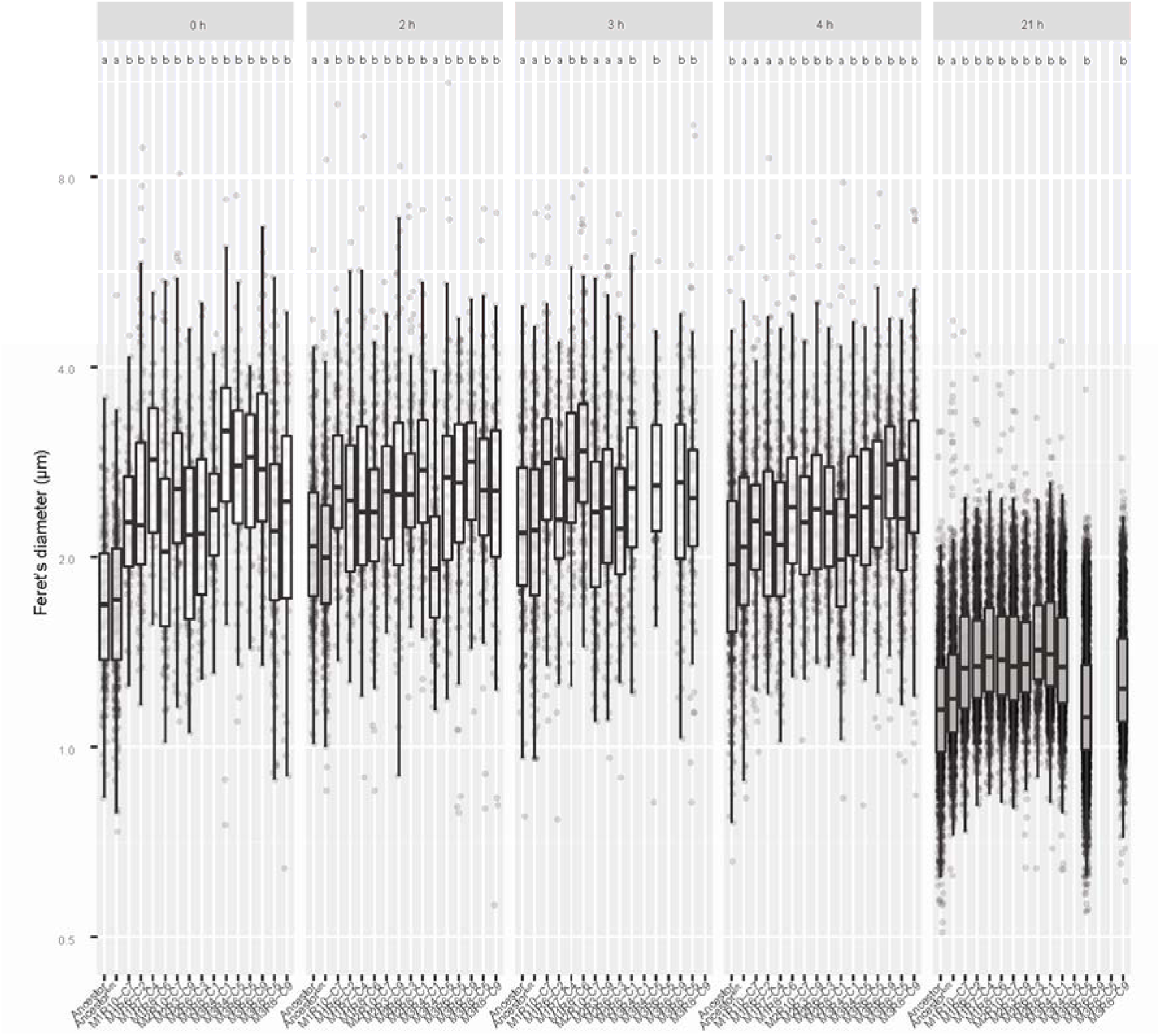
Size of individual *B. thetaiotaomicron* cells at different phases of growth. Cell size of ancestor, ancestor^tetR^and evolved M1-M3 lineages, shown by Feret’s diameter, which is the longest distance between any two points along the circumference of a bacterium, as determined with ImageJ2/Fiji (NIH). Every dot represents the diameter of a single bacterium. Evolved lineages were pregrown for 6 hours in the respective medium they were evolved in, while the ancestors were pregrown in BHIs. Time series indicates sizes of bacteria after dilution in fresh M3 medium. 0 h – 4 h are representative of logarithmic growth phase, after 21 h bacteria are in stationary phase. Significant differences (p<0.05) of evolved lineages or ancestor compared to ancestor^tetR^ are indicated by different letters. Significant differences between ancestor and evolved lineages or between evolved lineages are not depicted. Kruskal-Wallis rank sum test was used for overall comparison between different groups and the FDR method was used for pairwise comparison.

### Evolved lineages increase in cell size

Bacterial fitness is closely related to cell size. In E. coli, increase in cell size confers a FA [53, 54]. To determine whether the fitter evolved lineages also show an increase in cell size compared to the ancestor strain, we measured cell size during growth in M3 medium. Indeed, we observed an increase in cell diameter for all six M3-evolved lineages tested at almost all time points measured (p<0.05; Fig. 5 and Suppl. Fig. 4). This phenomenon is most pronounced during re-inoculation and logarithmic growth phase, with cell sizes ranging from a median of 2.2 to 3.2 µm for M3-evolved lineages at inoculation compared with a median of 1.7 µm for the ancestor. For M1- and M2-evolved lineages cell size increase during growth in M3 medium was less pronounced, but for most measurements median cell diameter was also significantly larger compared to the ancestor (p<0.05; Fig. 5).

### M3-evolved lineages deplete glucose more rapidly than the ancestor

To identify further factors contributing to the FA of evolved populations and lineages over the ancestor strain, selected evolved populations and lineages as well as the ancestor were subjected to glucose depletion experiments. Three out of four M3-evolved lineages had faster glucose depletion rates compared to the ancestor when grown in M1 medium (Suppl. Fig. 5a). Also, higher optical densities were reached after 5 h for the respective evolved lineages, indicating faster growth rates (Suppl. Fig. 5a). For M3-evolved lineages, glucose depletion in M3 medium (Suppl. Fig. 5b) was faster than during competition (Suppl. Fig. 5d). Concomitantly performed competition experiments also show an increase in FA of the lineages over the ancestor (Suppl. Fig. 5c). An initial increase in glucose, most probably due to amylopectin degradation and liberation of free glucose, occurred earlier in the M3-evolved lineages as compared to those lineages competed with the ancestor (Suppl. Fig. 5b, d). The peak of glucose in the ancestor alone was only present after 5 h, suggesting a late adaption to the new medium (Suppl. Fig. 5d). Furthermore, an inverse relationship between glucose depletion (at 6 hours) and FA (at endpoint) was observed, indicating a link between glucose consumption and increase in fitness, which might be directly correlated to growth rates (Suppl. Fig. 5e, f). As mentioned above, competition with the evolved lineages compromised growth of the ancestor beginning at 3 up to 5 h (Suppl. Fig. 5g).

### Early activation of starch utilization gene transcription in M3-evolved populations

The repression of complex glycan degradation PULs in the presence of simpler carbohydrates, also referred to as nutrient prioritization, has been reported in *B. thetaiotaomicron* [55, 56]. Gene expression analysis of M3-evolved lineages revealed that the *susC^starch^* (PUL66, encoding the TBDT for starch, amylopectin, amylose, maltooligosaccharides, and maltose transport) has higher expression levels in the presence of amylopectin regardless of the presence of glucose (Fig. 6a). Growth of M3- evolved lineages in the presence of both amylopectin and glucose revealed elevated mRNA transcripts of *susC^starch^* within 30 min and up to 3 h (Fig. 6a). Only after 4 h, when glucose was mostly depleted from the medium (Fig. 6c), did the ancestor increase *susC^starch^* expression. *susD^starch^*, which encodes an extracytoplasmic lipoprotein necessary for starch binding, followed similar expression kinetics although with less pronounced differences (Fig. 6b). A further indicator of highly active amylopectin degradation is that glucose levels increase within 30 min of M3-evolved lineages being exposed to amylopectin-containing media (Fig. 6c). This extensive release of glucose is not present in the ancestor, suggesting slower amylopectin degradation. Collectively this data suggest that M3-evolved lineages show increased Sus^starch^ operon expression in the presence of glucose, allowing for immediate degradation of amylopectin concomitant with glucose utilization rather than nutrient prioritization and a diauxic metabolic shift.

**Fig. 6.**
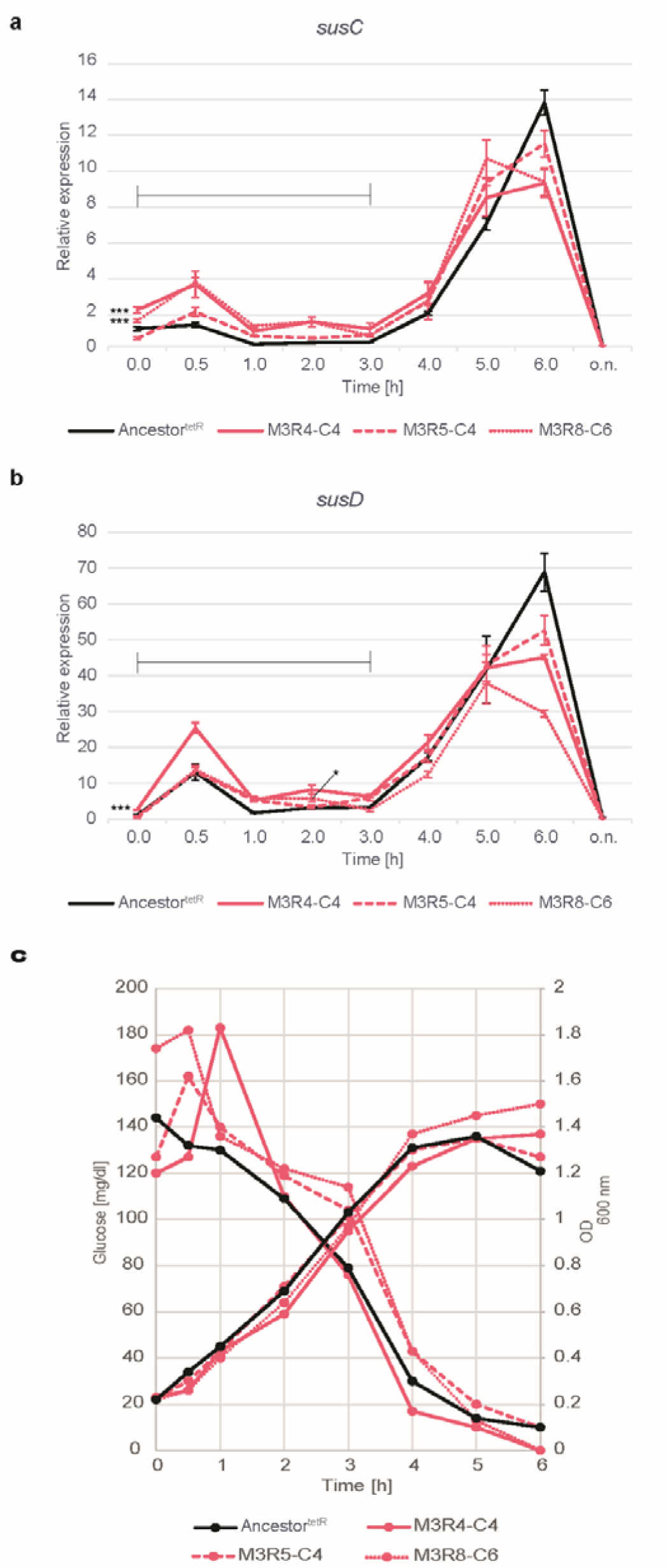
Transcriptional regulation of susC and susD. Relative expression of *susC^starch^* (**a**) and *susD^starch^* (**b**)of ancestor^tetR^ (black lines) and three M3-evolved lineages (pink lines) upon transfer to new medium with glucose and amylopectin, determined with RT-qPCR. The primer sequences can be found in Suppl. Table 5. Error bars indicate standard deviation from technical triplicates. (c) Glucose release and depletion as well as optical density OD_600 nm_ values upon transfer to new medium with glucose and amylopectin. Significant differences in relative expression between 0-3 h (pointed out by black horizontal lines) were determined by calculating the area under the curve for every lineage and the ancestor, followed by an ANOVA with Tukey’s HSD to correct for multiple comparisons (*p-value≤0.05, ***p-value≤0.001).

## Discussion

Bacteria can evolve rapidly in laboratory conditions and animal models with considerable parallelism across independent replicates [51, 57, 58]. Here we show that genetic diversification of *B. thetaiotaomicron* can occur within short timescales and is driven in part by the complexity of the nutrient landscape. M2- and M3-evolved populations showed a higher genetic diversification compared to M1-evolved populations, indicating a distinction between evolution driven by growth on a single monosaccharide (glucose) versus polysaccharides (Fig. 3). The increased range of available nutrients might provide *B. thetaiotaomicron* with the ability to thrive in additional niches due to a higher prevalence of possible novel traits, as has been suggested for E. coli evolving in media with different complexities of antibiotics [59].

*B. thetaiotaomicron’s* rapid genetic diversification was characterized by SNP and INDEL accumulation as well as structural variants (shufflons). Phase variation is often connected to pathogenicity and bacteriophage defense [60, 61]. In the human gut, phase variation may also help bacteria to withstand perturbations (e.g. variations in host diet) by creating phenotypic heterogeneity [62]. We identified both characterized as well as novel phase variation loci, three of which involve SusC-like proteins. The SusC-like proteins in the first gene cluster are a part of PUL14, which is important for complex N-glycan and O-glycan uptake [5, 27] (Suppl. File 1 PUL table). The second gene cluster involves SusC-like proteins within a predicted peptide-binding PUL (BT2264) [17] that are upregulated in *B. thetaiotaomicron* mono-colonized transgenic colitis rats (BT2260) [63] as well as in IgA-expressing mice (BT2268, Mucus-Associated Functional Factor, MAFF) [64] , suggesting potentially important functions in mucus colonization and degradation. Future investigations are needed to elucidate what factors affect shufflon activity levels and their phenotypic consequences.

Long-term evolution experiments in E. coli have demonstrated increased competitive fitness that is accompanied by the accumulation of mutations throughout 50,000 generations [65]. For *B. thetaiotaomicron*, we observed a higher FA in more complex competition media and when populations have evolved in a complex medium (Fig. 4). The fitness increase of populations and lineages indicates the presence of adaptive genetic diversification. The FA of evolved lineages was highly variable compared to evolved populations, with several lineages having a very low FA while others outcompeted the ancestor completely. This indicates that mutational profiles of single lineages vary and that differences in the mutation profile in the evolved populations augment the fitness of the population in comparison to certain clonal competitors. We suggest that FA is a complex trait involving multiple mutations and different combinations of mutations. Population-level FA may also involve inter-lineage interactions. Whole genome sequencing of many evolved lineages, as well as reverse genetics approaches, will be needed to further decipher the specific mutations that contribute to the FA of the evolved lineages and populations.

Several factors may contribute to the observed FA. Evolved populations and lineages of *B. thetaiotaomicron* developed physiological adaptations that include an increase in cell size during growth, the ability to inhibit the growth of the ancestor, and efficient substrate use. A direct correlation of bacterial cell size with growth rate has been postulated for several bacterial species [54, 66]. Cell volume increases were also found for *E. coli* in a long-term evolution experiment [54]. We found an increase in the size of *B. thetaiotaomicron* evolved in M1-, M2- and M3 medium upon growth in the complex M3 medium in comparison to the ancestor (Fig. 5, Suppl. Fig. 4). One advantage of a larger cell is the possibility to accommodate more DNA and thereby facilitate multifork replication [67]. This allows the cell to proceed with DNA replication before cell division, thereby leading to faster growth [66]. Whether such a mechanism is present in fast-growing *B. thetaiotaomicron* needs still to be determined.

Prioritization of certain glycan substrates has been described for *B. thetaiotaomicron* [68]. Preferred carbohydrate substrates lead to the repression of PULs for lower-priority carbohydrates, which can lead to diauxic growth. Besides substrate prioritization, synergistic substrate use is possible in *B. thetaiotaomicron* [56, 68]. Whether PULs are repressed or not is dependent on the substrate combinations. In the highly complex gut environment, with fluctuating substrate concentrations, these regulations might be pivotal in guaranteeing the persistence of *B. thetaiotaomicron*. Canonical SusC, within PUL66, responsible for starch and amylopectin import into the periplasm, is not repressed in the presence of glucose [68]. Exposing evolved lineages to medium with both amylopectin and glucose increased relative expression of susC and susD transcript levels in comparison to the ancestor^tetR^ (Fig. 6). A faster response in sensing glycans, i.e. amylopectin, and upregulation of respective PUL machinery to import and catabolize such might enable *B. thetaiotaomicron* evolved populations and lineages to gain higher FA compared to ancestor^tetR^.

In the intestinal tract, rapid adaptive evolution, driven in part by fluctuating nutrient availability and adaptation to the dietary patterns of the host, may be one mechanism that supports the long-term stability of an individual person’s microbiome. Whether microbial competition for substrates with other species accelerates the process of adaptive evolution still needs to be investigated. Future work should evaluate in vitro and in vivo evolution in multi-species consortia to decipher the influence of other species on *B. thetaiotaomicron* evolution.

## List of Abbreviations

BHIs,: brain heart infusion medium with supplements;
CAZymes,: carbohydrate-active enzymes;
CLSM,: confocal laser scanning microscope;
ECF-type sigma factors,: extracytoplasmic function-type sigma factors;
FA,: fitness advantage;
HTCS,: hybrid two-component system
MFS,: major facilitator superfamily;
MM,: minimal medium;
OD,: optical density;
PUL,: polysaccharide utilization loci;
RGI,: rhamnogalacturonan I;
rpoB,: RNA polymerase beta;
SNP,: single nucleotide polymorphism;
Sus,: starch utilization system;
TBDT,: TonB-dependent transporter;
TetR,: tetracycline resistance;
XOS,: xylooligosaccharides;

## Data availability

Sequencing data have been deposited at the National Center for Biotechnology Information (NCBI) Sequence Read Archive under the BioProject number PRJNA1257038 (https://dataview.ncbi.nlm.nih.gov/object/PRJNA1257038?reviewer=tusv2tutm20il769netoq4ks9l). Scripts and pipelines relevant to the manuscript are deposited at github https://github.com/vakilos/bacevo.

## Author Contributions

M.L. designed and conducted experiments, acquired, analyzed and interpreted the data and wrote the manuscript, C.Z. conducted experiments, acquired, analyzed and interpreted the data, developed analytical tools; A. H., N.I., J. S. and K. J. F. conducted experiments and analyzed data. O.B. analyzed and interpreted the data, developed analytical tools; F. C. P. acquired and analyzed data; S.K. analyzed and interpreted data, D.B. conceived the study, designed experiments, analyzed and interpreted the data, supervised the project, and critically revised the manuscript. All authors read and approved the manuscript.

## Acknowledgements

We want to thank Nicole Nussbaum and Georgi Nikolov for technical assistance. We thank Eric Martens (University of Michigan Medical School, USA) for kindly providing the Bacteroides species integrative vector pNBU2-bla-tetQb (Tet^R^, Amp^R^).

The metagenome sequencing was performed by the Next Generation Sequencing Facility at the Vienna BioCenter Core Facilities (VBCF), member of the Vienna BioCenter (VBC), Austria. Whole genome sequencing of evolved lineages was performed by the Joint Microbiome Facility (JMF) of the University of Vienna and the Medical University of Vienna.

## Supplementary figure legend

**Suppl. Fig. 1 Distribution of mutations among PUL genes**

**a** Number of mutations per nucleotide in PUL genes: carbohydrate esterases (CE), polysaccharide lyases (PL), sulfatases, hybrid two-component systems (HTCS), SusC-like proteins (SusC), glycosyl hydrolases (GH), SusD-like proteins (SusD), ECF-type sigma/anti-sigma factors, and all genes in the entire genome of each evolved population (grey dots) evolved in different media. Kruskal-Wallis rank sum test with Dunn’s correction was used to reveal any significant changes between M1-M3 evolved populations (n_M1-evolved_=5, n_M2-evolved_=7, n_M3-evolved_=6) for any PUL gene or all genes. *p-value <0.05, **p- value <0.01 **b** Number of mutations per nucleotide in PUL genes and all genes independent of the media the population replicates have evolved in. Significant differences between groups of genes were analyzed using Aligned Ranks Transformation ANOVA with Tukey’s post-hoc correction and are indicated by different letters.

**Suppl. Fig. 2 OD_600nm_ values of evolved populations and lineages during competition**

Violin plot of OD_600nm_ values of the (**a, b**) ancestor and M1-M3-evolved populations and (**c, d**) lineages in stationary phase at the end of the competition against ancestor^tetR^. Every dot represents the OD value of an evolved population or the ancestor competed against ancestor^tetR^. Significant differences of OD_600nm_ values between the two groups were analyzed using Kruskal-Wallis test and are indicated by different letters. OD_600nm_ values of ancestor and M1-M3-evolved populations competed in (**a**) BHIs, M1 –M3 medium (G, glucose; AP, amylopectin; AG, arabinogalactan; I, inulin; P, pectin; XOS, xylooligosaccharides), or (**b**) with single glycans of the complex M3 medium in MM. OD_600nm_ values of M1-M3-evolved lineages competed in (**c**) BHIs, M1–M3 medium, or (**d**) with single glycans of the complex M3 medium in MM. Colours indicate different lineages from various replicates.

**Suppl. Fig. 3 Compromised ancestor growth**

**a** Number of generations of ancestor^tetR^, ancestor, and M3-evolved lineages during competition and without competition over time. Ancestors and M3-evolved lineages were grown for 21 hours in fresh M3 medium with or without competition. After an initial exponential growth of both the ancestor and the evolved lineages, the ancestor stalled in growth or even dropped in cell numbers. **b** Percentage of compromised or dying cells during competition (yellow border) and without competition after 4 hours of growth. Error bars indicate standard deviation from 3-9 images analyzed each condition. Statistical significant differences between growth alone (no competition) and with competition were calculated using Independent Samples T-test (*p<0.05, **p<0.01). **c-f** Lookup Table (LUT) images from DAPI stained *B. thetaiotaomicron* cells as presented by the Leica Application Suite X software from images taken on a confocal laser scanning microscope. Compromised cells with low DNA content (presented in black in the LUT image) are highlighted by a yellow arrow. LUT images of ancestor^tetR^ competed with ancestor or M3-evolved lineages after 4 hours of growth, at the peak of cell damage, (**c, e**) or of the ancestors and lineages alone (**d, f**). The scale bar shown is 10 µm in length.

**Suppl. Fig. 4 Bacterial cell size images**

Representative DAPI images (in blue) of ancestor^tetR^ and ancestor and M3-evolved lineages grown for 0-21 h in fresh M3 medium, after pregrowth for 6 h in M3 medium for evolved populations and BHIs medium for ancestors. For better visualization, all images have been cropped to the same extent. The scale bar shown is 10 µm in length.

**Suppl. Fig.5 Glucose levels and growth conditions in M1 and M3 medium**

**a** Glucose depletion and OD increase in M3R5, four M3-evolved lineages, and ancestor^tetR^ over 5 h in M1 medium. M3 evolved-population and – lineages were pregrown in M3 medium, ancestor^tetR^ in BHIs. Glucose depletion between the ancestor^tetR^ and M3-evolved lineages was calculated using Friedman rank sum test and Conover’s single-step correction. (**b**) Glucose depletion of four M3- evolved lineages and ancestor^tetR^ without competition. (**c**) FA of M3-evolved populations of the experiment performed in **d**. (**d**) Glucose depletion of four M3-evolved lineages and ancestor^tetR^ during competition for 21 h. Correlation between FA at 21 h and glucose levels after 6 h without competition (**e**) and during competition (**f**). Glucose levels negatively correlate with FA with a Pearson’s correlation coefficient r=0.99 and r=0.52 without competition and during competition, respectively. (**g**) Number of generations of ancestor^tetR^, ancestor, and M3-evolved lineages during competition and without competition over time. Ancestor^tetR^ and M3-evolved lineages were grown for 21 hours in fresh M3 medium with or without competition.

**Suppl. Fig. 6 Construction of a *B. thetaiotaomicron^tetR^* derivative strain**

**a** Schematic representation of the pNbu2-bla-tetQb plasmid and recombination events leading to its integration into one of the two *B. thetaiotaomicron* possible tRNA^ser^ loci, *attN2-1* and *attN2-2*. Primers used to verify insertion of the plasmid in either of the loci (Nbu2-*att1* FW and Rev for locus 1 and Nbu2-*att2* FW and Rev for locus 2) are shown, as well as the expected size of amplified genome fragments flanked by each primer pair prior to plasmid integration. Features of the plasmid shown: origin of replication (*oriR*), integrase gene (*intN2*), β-lactamase gene for ampicillin selection in *E. coli* (*bla*), tetracycline gene for selection in Bacteroides spp. (*tetQ*) and recombination site (*attN2*). **b** Growth, measured as OD , of **B. thetaiotaomicron** wild type (WT) and Tet^R^ transconjugant strain in BHIs liquid medium (orange line) or in BHIs medium supplemented with 5 μg ml^-1^ erythromycin (Erm; blue line) or 3 μg ml^-1^ tetracycline (Tet; purple line). Pictures of the culture vials at the end of experiment are shown for each strain (right panel). **c** Chromosomal DNA of Tet^R^ (and Erm^R^, not used in this experiment) **B. thetaiotaomicron** conjugants were screened by PCR using primer pairs Nbu2-*att1* FW/Rev and Nbu2-*att2* FW/Rev to confirm insertion of the *pNbu2-bla-tetQb* plasmid. DNA amplification with *att1* primers yields a band with high molecular weight (> 6,000 bp) for the Tet^R^ strain (referred to as ancestor^tetR^) that contrasts with a 729 bp band obtained for the WT strain, confirming plasmid insertion at locus 1. L: DNA ladder; MW: molecular weight, in base pairs (bp). *Unspecific amplification product.

**Suppl. Fig. 7 Effect of pregrowth conditions of the ancestor on FA**

Paired Wilcoxon signed-rank test was used to compare FA of M3-evolved populations or ancestor and ancestor^tetR^, under different pregrowth conditions. The ancestor and ancestor^tetR^ were pregrown in either BHIs or M3 medium, M3-evolved populations in M3 medium. Competition experiments were performed. Ancestor pregrowth in M3 medium significantly reduced the FA of the M3-evolved populations by a factor of 2 (p=0.036), during competition in M3 medium.

