## Supplementary figures and images for "Nutrient landscape shapes the genetic diversification of the human gut commensal *Bacteroides thetaiotaomicron*"

### Supplementary Figure 1

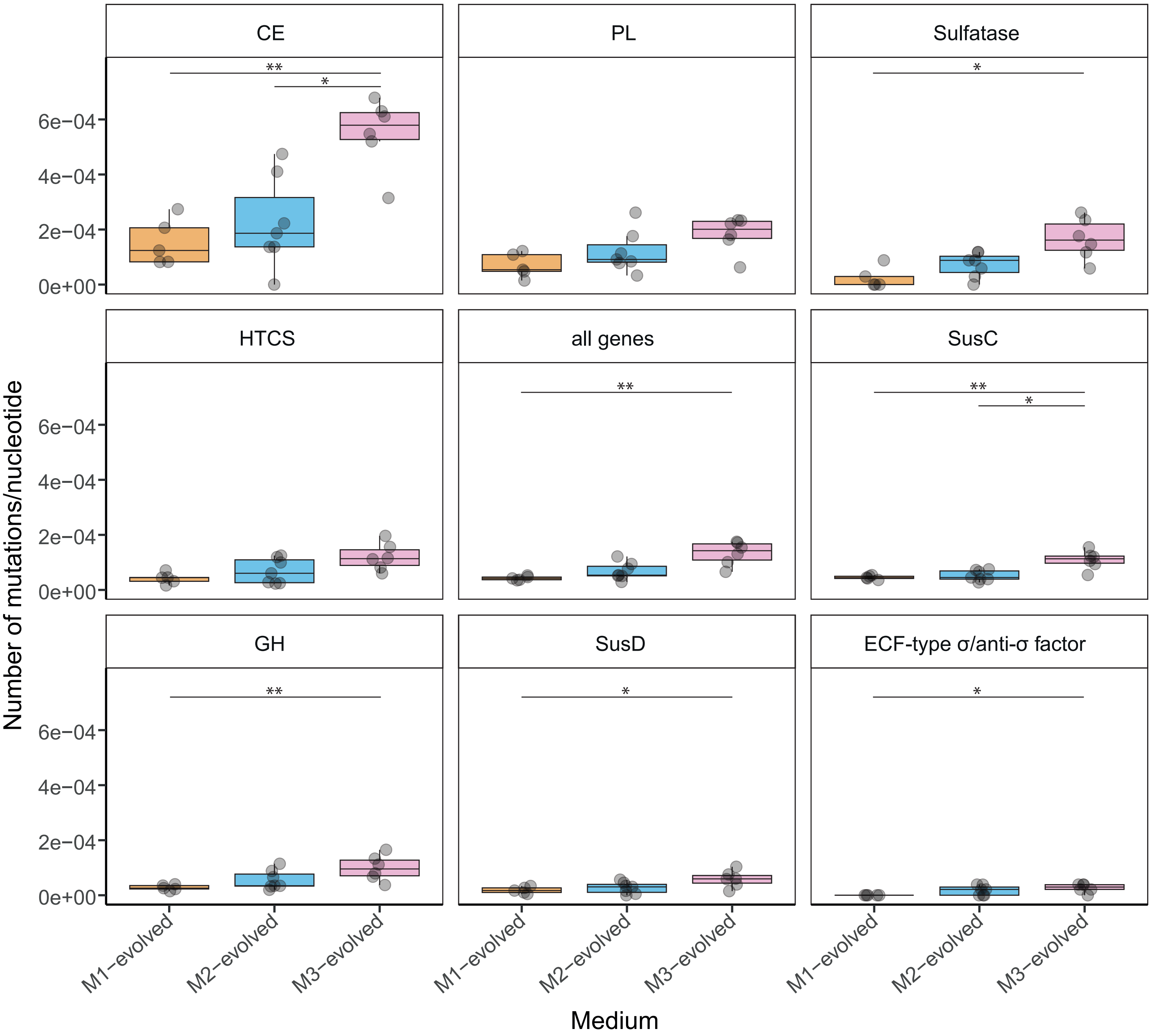

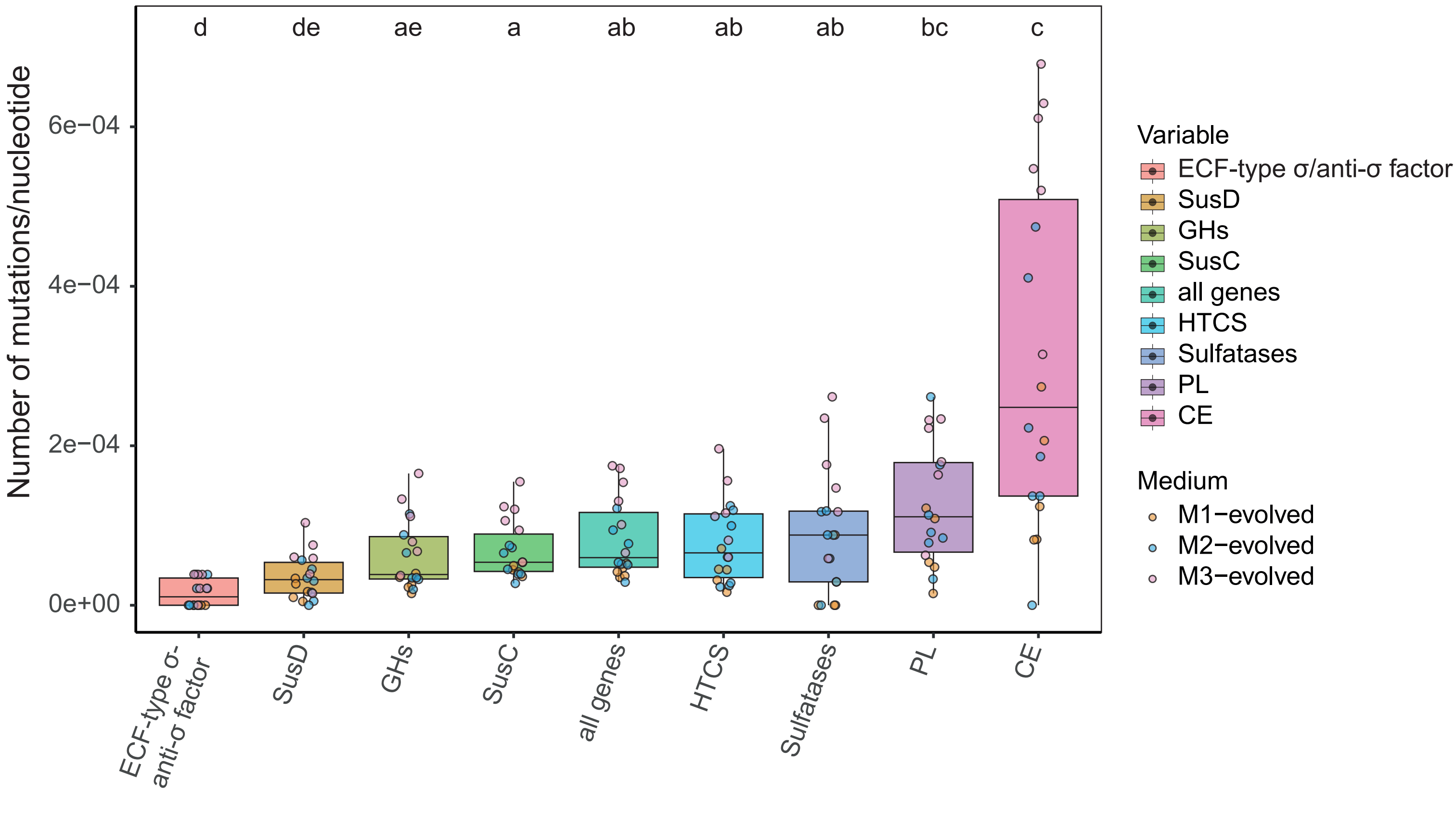

### Supplementary Figure 2

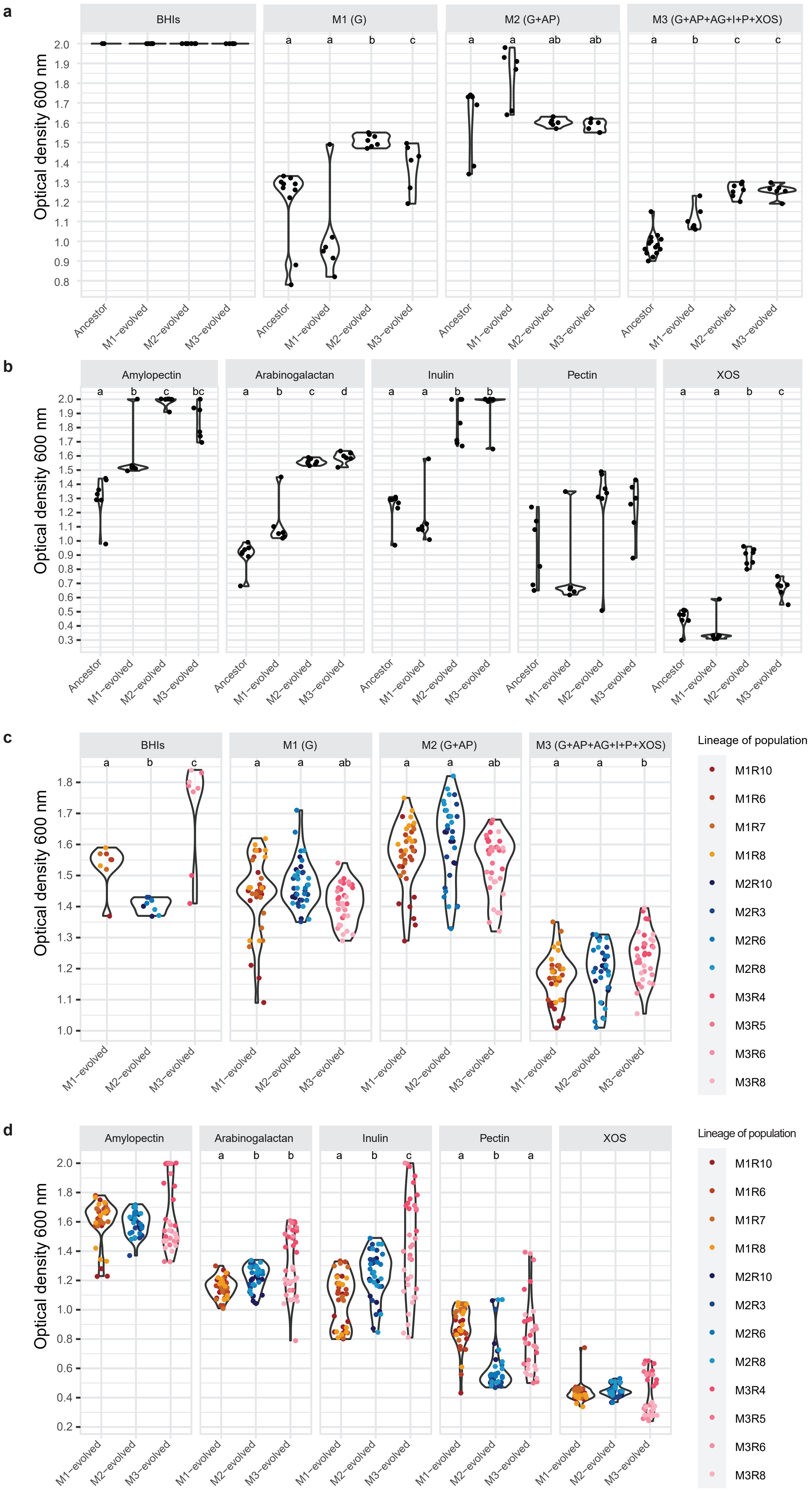

### Supplementary Figure 3

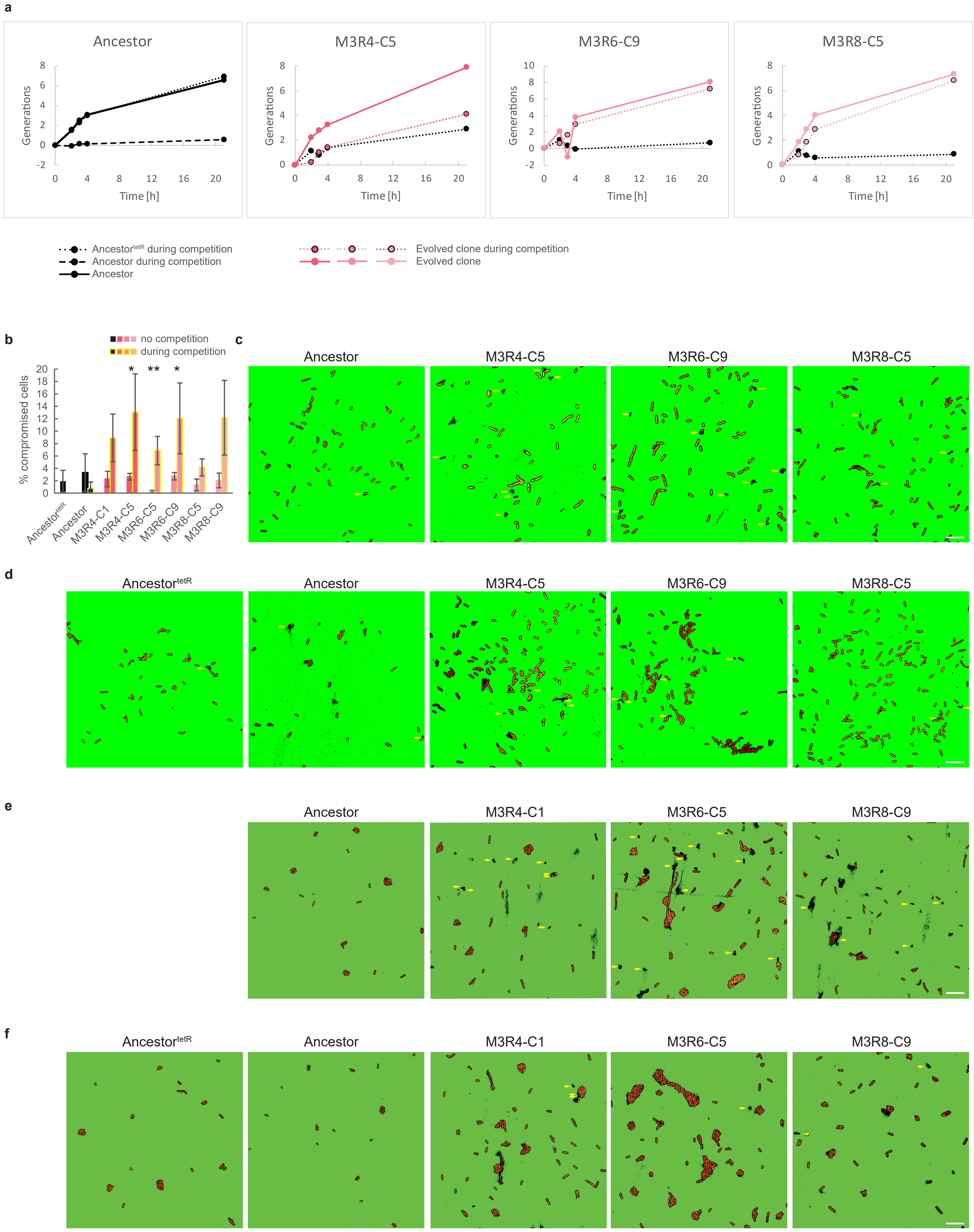

### Supplementary Figure 4

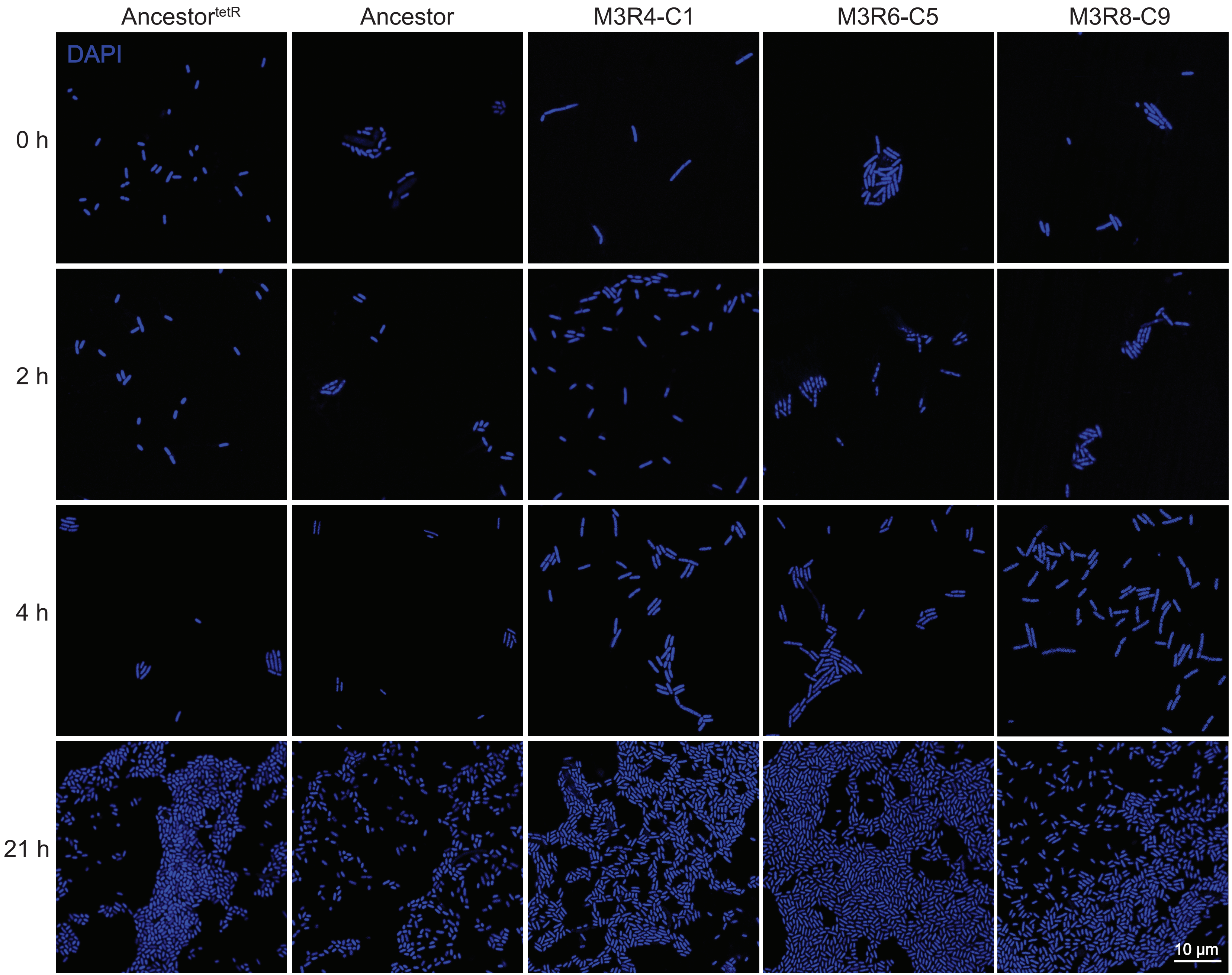

### Supplementary Figure 5

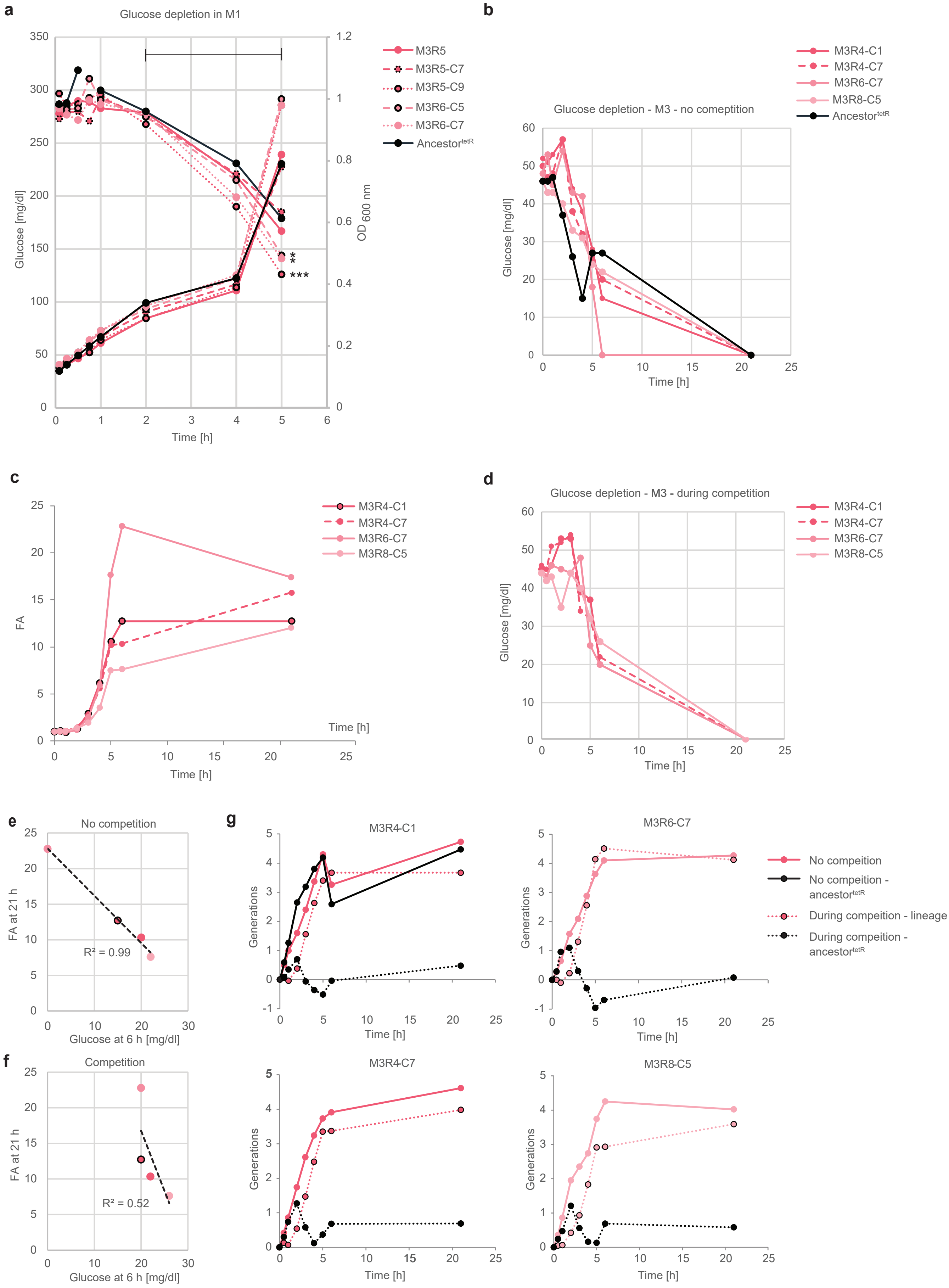

### Supplementary Figure 6

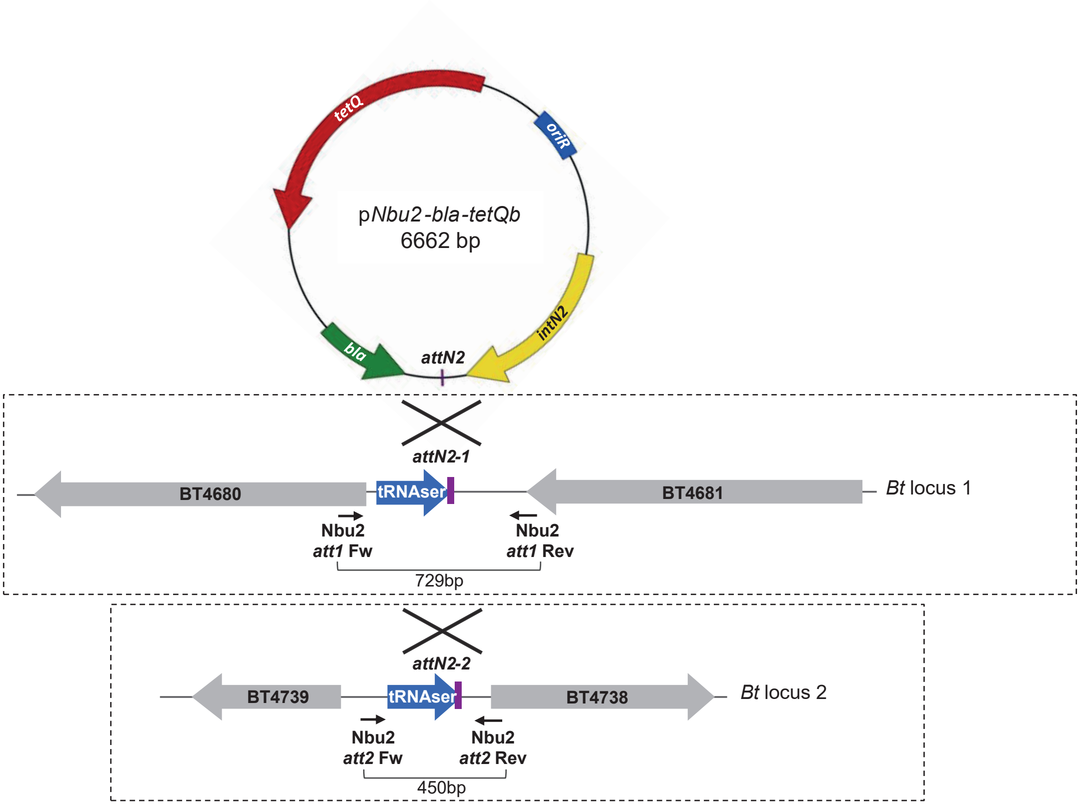

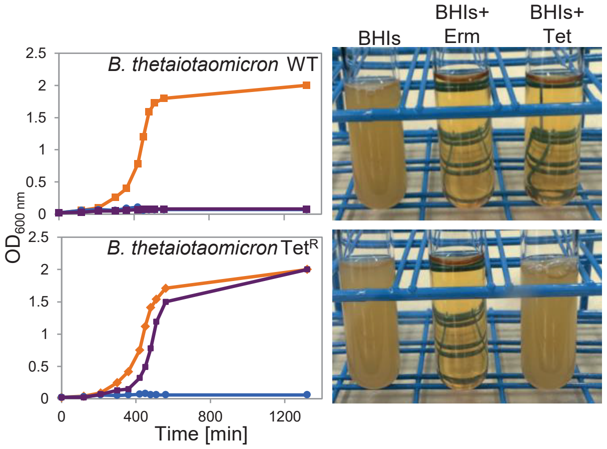

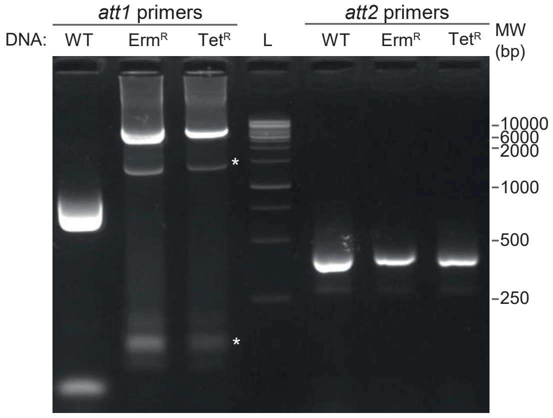

### Supplementary Figure 7

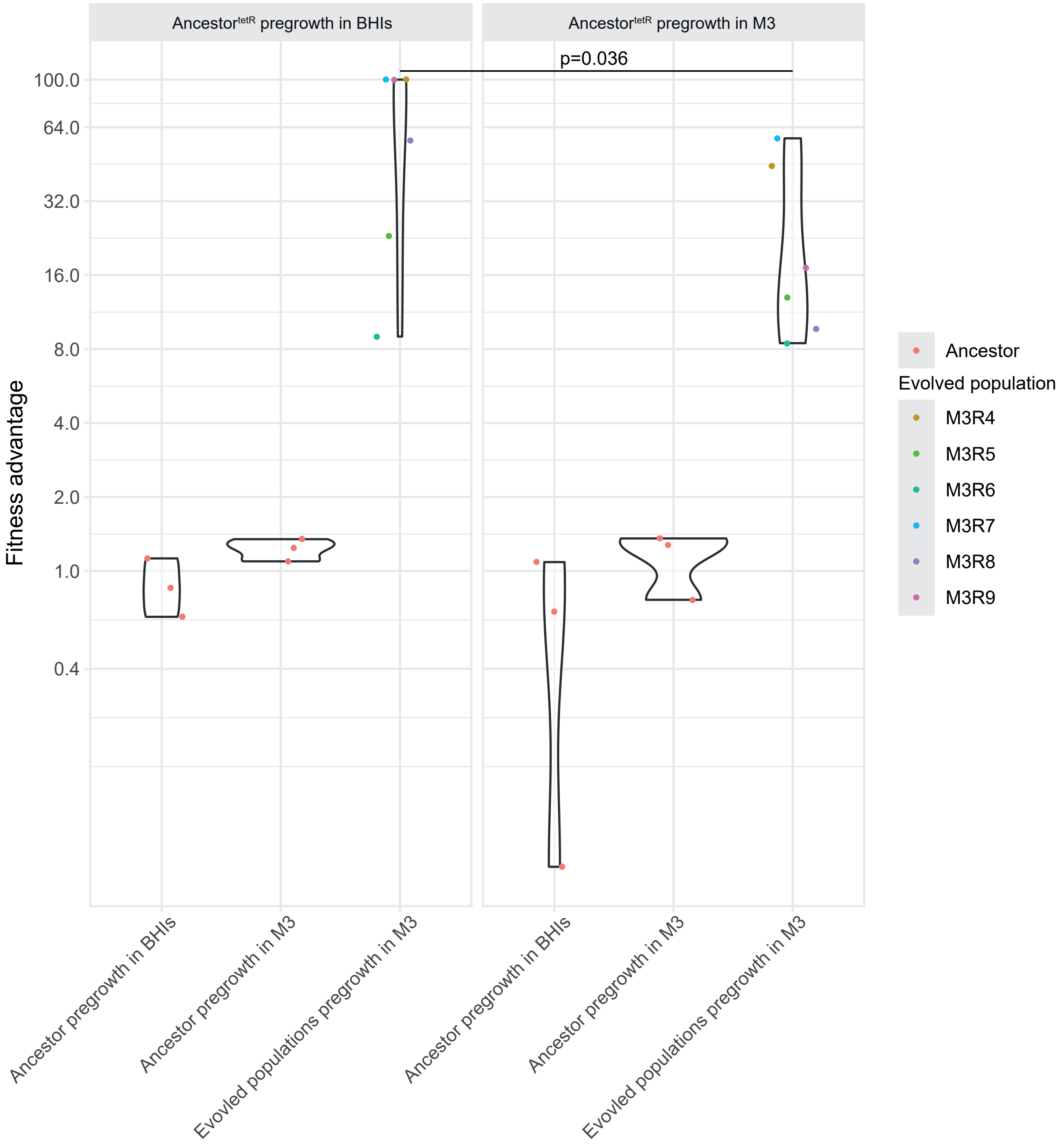
