## Supplementary Information for "Nutrient landscape shapes the genetic diversification of the human gut commensal *Bacteroides thetaiotaomicron*"

**Supplementary material**

**Supplementary tables**

**Supplementary table 1. Minimal medium composition**

| **Medium component** | **Company** | **Article number** | **Molecular weight [g/mol]** | **g/l medium** | **Final concentration [mM]** |
| --- | --- | --- | --- | --- | --- |
| **KH_2_PO_4_**, ≥99 %, p.a., ACS | Carl Roth | 3904 | 136.09 | 0.9 | 6.60 |
| **NaCl**, ≥99.5%, BioUltra, for molecular biology | Sigma-Aldrich | 71376 | 58.44 | 0.9 | 15.40 |
| **MgCl_2_·6H_2_O**, ≥99 %, BioXtra | Sigma-Aldrich | M2670 | 203.3 | 0.02 | 0.098 |
| **CaCl_2_·2H_2_O**, ≥99 %, BioXtra | Sigma-Aldrich | C5080 | 147.01 | 0.026 | 0.177 |
| **CoCl_2_·6H_2_O**, BioReagent | Sigma-Aldrich | C8661 | 237.93 | 0.001 | 0.0042 |
| **MnCl_2_·4H_2_O**, ≥99 %, p.a | Carl Roth | T881 | 197.91 | 0.01 | 0.0505 |
| **NH_4_Cl**, ≥99.5%, for molecular biology | Sigma-Aldrich | A9434 | 53.49 | 0.5 | 9.35 |
| **Na_2_SO_4_**, ≥99 %, p.a., ACS | Carl Roth | 8560 | 142.04 | 0.25 | 1.76 |
| ***NaHCO_3_**, ≥99,5 %, CELLPURE^®^ | Carl Roth | HN01 | 84.01 | 2 | 23.81 |
| **9.99 mM FeSO_4_·7H_2_O**, ≥99 %, p.a., ACS | Carl Roth | P015 | 278.01 | 1.5 ml | 0.01499 |
| **L-cysteine**, ≥98 %, DAB, for biochemistry | Carl Roth | 3467 | 121.16 | 1 | 8.25 |
| ***Vitamin B12**, ≥96 %, for biochemistry | Carl Roth | T915 | 1355.37 | 0.0001 | 7.38E-05 |
| ***Vitamin K1**,≥97 %, for biochemistry | Carl Roth | 3804 | 450.7 | 0.000001 | 2.22E-06 |
| ***Hemin** from porcine, ≥98 %, BioXtra | Sigma-Aldrich | 51280 | 651.94 | 0.005 | 7.669 |
| ***Resazurin sodium salt**, BioReagent | Sigma-Aldrich | R7017 | 251.17 | 0.001 | 0.0039 |

*these components were added after autoclaving under sterile conditions

**Supplementary table 2. Detailed information on carbohydrates**

| **Carbohydrate** | **Company** | **Article number** |
| --- | --- | --- |
| Amylopectin from maize | Sigma-Aldrich | 10120 |
| Arabinogalactan from larch wood | Sigma-Aldrich | 10830 |
| D(+)-Glucose, p.a., ACS, anhydrous | Carl Roth | X997 |
| Inulin from chicory | Sigma-Aldrich | I2255 |
| Pectin from apple | Sigma-Aldrich | 76282 |
| Xylooligosaccharides from corncob, ≥95 % | Carl Roth | 8659 |

**Supplementary table 3. BHI medium composition**

| **Medium component** | **Company** | **Article number** | **g/l medium** |
| --- | --- | --- | --- |
| **Brain heart infusion broth**, Oxoid™ | ThermoFisher Scientific | CM1135 | 37 |
| **Yeast extract**, Bacto™ | ThermoFisher Scientific | 212750 | 5 |
| ***NaHCO_3_**, ≥99,5 %, CELLPURE^®^ | Carl Roth | HN01 | 1 |
| **L-cysteine**, ≥98 %, DAB, for biochemistry | Carl Roth | 3467 | 1 |
| ***Hemin** from porcine, ≥98 %, BioXtra | Sigma-Aldrich | 51280 | 0.005 |
| ***Vitamin K1**, ≥97 %, for biochemistry | Carl Roth | 3804 | 0.001 |

*these components were added after autoclaving under sterile conditions

**Supplementary table 4. PCR primers to confirm pNUB2-bla-tetQb integration into *B. thetaiotaomicron***

| **Primer name** | **Primer sequence (5'->3')** | **Ref.** |
| --- | --- | --- |
| Nbu2-att1 Fw | CCTTTGCACCGCTTTCAACG | ^1^Martens et al. |
| Nbu2-att1 Rev | TCAACTAAACATGAGATACTAGC |  |
| Nbu2-att2 Fw | TATCCTATTCTTTAGAGCGCAC | ^1^Martens et al. |
| Nbu2-att2 Rev | GGTGTACCTGGCATTGAAGG |  |

**Supplementary table 5. qPCR primers**

| **Primer name** | **Primer sequence (5'->3')** | **Length** | **GC%** | **Gene** | **Gene name** | **Ref.** |
| --- | --- | --- | --- | --- | --- | --- |
| rpoB-Fw | CAAATCGGACGCAAGTCAATAG | 22 | 45.5 | DNA-directed RNA polymerase subunit β | BT2734 | This work |
| rpoB-Rev | CCGTACAGAGGACGAAGAATTT | 22 | 45.5 |  |  |  |
| tetR-Fw | AGAGCATCGGTTCGAGAATG | 20 | 50 | TetQ, Tetracycline resistance | TetQ | This work |
| tetR-Rev | CGGTAAGTACACCTGCTGATT | 21 | 47.6 |  |  |  |
| 16S-FW | CGTTCCATTAGGCAGTTGGT | 20 | 50 | 16S |  | ^2^Whitaker et al. |
| 16S-Rev | CAACCCATAGGGCAGTCATC | 20 | 55 |  |  |  |
| SusC-Fw | CGGTAAAATAGCCGGTGTCA | 20 | 50 | starch utilization system C | BT3702 | ^3^Liu et al. |
| SusC-Rev | GTTTCGATATCCGCAGGGTT | 20 | 50 |  |  |  |
| SusD-Fw | CAATCTCGAAGATGTGCCGA | 20 | 50 | starch utilization system D | BT3701 | ^3^Liu et al. |
| SusD-Rev | GTCTGAATTTTGCGTCTGCC | 20 | 50 |  |  |  |

**Supplementary results**

**Mutations in specific PULs involved in complex glycan degradation**

To verify an increasing number of mutations in specific genes involved in degradation of carbohydrates used in the study we investigated such in more detail. For arabinogalactan degradation two PUL are known (PUL5, BT0262-BT0291 and PUL65, BT3674-BT3687)^4-7^. Within PUL5 both SusC homologues BT0268 and BT0272 show more mutations in M3-evolved populations compared to M1- or M2-evolved populations, whereas for BT0272 only in M3-evolved populations mutations are present.

XOS have a degree of polymerization between 2 and 10 and a maximum of up to 20 xylose units linked by β-(1-4) glycosidic bonds^8^. XOS are derived from xylans. Depending on the source of xylan, the degree of acetylation can vary. For corn cob xylan is has been shown that it contains about 0.6 % (w/w) of acetyl content or 3.3 [mol%], suggesting XOS acetylation. Acetylation can take please at O-2 and/or O-3^8^ (Fig. 1b). Acetyl xylan esterases (EC3.1.1.72) catalyze the hydrolysis of the acetyl group from XOS. *B. thetaiotaomicron* has two putative acetyl xylan esterases, BT2525, belonging to the carbohydrate esterase family 7 (CE7) and BT3246 (CE4). Whereas no mutations have been found in BT2525, BT3246 shows an increasing number of SNP in M3-evolved populations compared to M1- and M2-evolved populations. Furthermore, the acetyl xylan esterases BT3246 is the only gene mutated within the putative PUL50 with unknown substrate, suggesting a specific adaptation to M3 medium and xylan or XOS as a potential substrate for this PUL.

Pectin is by far the most complex carbohydrate used in the study and thus, 7 different PUL are known to be involved in the degradation of pectin so far. It consists of rhamnogalacturonan I (RGI), homogalacturonan, xylogalacturonan, and rhamogalacturonan II. In the periplasm, sides chains in RGI containing arabinosyl and galactosyl oligosaccharides need to be removed from the backbone before backbone depolymerization. This is orchestrated by three synergistically acting exo-β-1,4-galactosidases (GH2, EC3.2.1.23), BT4151, BT4156, and BT4160^9^. Within this highly complex PUL for RGI degradation, galactosidase BT4151 presented with a high number of low frequency mutations in M3-evolved populations compared to M1- and M2-evolved populations. Mutations in other genes within this PUL were rare. Another PUL involved in homogalacturonan and rhamogalacturonan II degratdation is PUL75^3,4,9,10^ (BT4108-BT4124). Number of low frequency mutations in M3-evolved populations increased in the HTCS response regulator BT4111 and in the SusD homologue BT4122. Mutations in other genes within this PUL such as the pectate lyase BT4118, which cleaves of (1-4) α-D-galacturonan, and the polygalacturonase BT4123 were similarly often mutated in M2-evolved populations. To conclude, *B. thetaiotaomicron* evolved populations gain mutations within specific genes important for complex carbohydrate degradation.

**Effect of pre-growth conditions on competition performance**

To examine the effect of pre-growth conditions on competition performance, *B. thetaiotaomicron^ancestor-tetR^* and *B. thetaiotaomicron^ancestor^* were pre-grown in BHIs and M3 medium and competed against each other in M3 medium. In addition, *B. thetaiotaomicron^ancestor-tetR^* pregrown in BHIs and M3 medium was competed in M3 against M3-evolved populations pre-grown in M3. Ancestor pre-growth in M3, compared to pre-growth in BHIs, increased FA by a factor of 2 (p=0.036; Suppl. Fig. 7). Pre-growth in the medium in which the competition is later on performed results in a slight advantage for the respective bacterial population. Compared to the effect of a 3-month evolution in such medium the effect is minor. Nevertheless, a certain effect of transcriptional adaption to the medium within hours might be present. Thus, a FA threshold of 2 was set to distinguish populations with a competition advantage from those without.
